# PolyGraph – Flexible, Biocompatible & Electrically Optimised Graphene-Polymer Composites for Next-Generation Neural Interfaces

**DOI:** 10.1101/2025.09.02.673516

**Authors:** Jack Maughan, Ian Woods, Cian O’Connor, Pablo Quintana-Sarti, Eoin Caffrey, Jose M. Munuera, Adrian Dervan, Alejandro López Valdés, Omar Mamad, Maeve A. Caldwell, Fergal J. O’Brien, Jonathan N. Coleman

**Affiliations:** School of Physics, Trinity College Dublin (TCD), Dublin, Ireland, D02 PN40; Tissue Engineering Research Group (TERG), Royal College of Surgeons in Ireland (RCSI), Dublin, Ireland, D02 YN77; Centre for Research on Adaptive Nanostructures and Nanodevices (CRANN) Trinity College Dublin (TCD), Dublin, Ireland, D02 W9K7; Advanced Materials and BioEngineering Research (AMBER) Centre RCSI and TCD, Dublin, Ireland, D02 W9K7; Department of Electronic and Electrical Engineering, School of Engineering Trinity College Dublin (TCD), Dublin, Ireland, D02 DP29; Trinity Centre for Biomedical Engineering (TCBE) Trinity College Dublin (TCD), Dublin, Ireland, D02 R590; Trinity College Institute of Neuroscience (TCIN), Trinity College Dublin (TCD), Dublin, Ireland, D02 PN40; Global Brain Health Institute (GBHI), Trinity College Dublin (TCD), Dublin, Ireland, D02 X9W9; Instituto de Ciencia y Tecnología del Carbono, INCAR-CSIC C/Francisco Pintado Fe 26, 33011 Oviedo, Spain; Department of Physiology & Medical Physics, Royal College of Surgeons in Ireland (RCSI), Dublin, Ireland, D02 YN77; FutureNeuro Research Ireland Centre for Translational Brain Science Royal College of Surgeons in Ireland (RCSI), Dublin, Ireland, D02 YN77; Discipline of Physiology, School of Medicine, Trinity College Dublin (TCD), Dublin, Ireland, D02 R590

**Author notes:** Funding: J. M., I. W., C. O’C., A. D., F. O’B and J. N. C. acknowledge the Research Ireland AMBER Centre for providing primary financial support to this study (SFI/12/RC/2278_P2). C. O’C. also acknowledges funding from an Anatomical Society Studentship. J. M. M., acknowledges funding from the Generation D initiative, promoted by Red.es, an organisation attached to the Ministry for Digital Transformation and the Civil Service, financed by the Recovery, Transformation and Resilience Plan through the European Union’s Next Generation funds.

**Keywords:** Graphene-polymer composites, flexible neural interfaces, biocompatible nanomaterials, electroactive biomaterials, neural stimulation, soft bioelectronics

## Abstract

Neural interfacing materials must deliver exceptional electrochemical performance, while integrating safely with the central nervous system. In this study we develop PolyGraph, a flexible, conductive, and biocompatible graphene-polycaprolactone (PCL) nanocomposite designed to strike this balance, which enables fabrication of conformable multichannel microelectrode arrays. Optimised liquid-phase exfoliation produces conductive, biocompatible PVP-stabilised graphene nanosheets, which are incorporated into PCL to form flexible, processable composites – PolyGraph. This material demonstrates bio- and immuno-compatibility with sensitive primary and iPSC-derived neuronal and glial cells. PolyGraph achieves low impedance (∼1.6 Ω cm^2^ @ 1 kHz) and high charge injection capacity (11.7 mC/cm^2^ for a 100 ms pulse), enhanced by NaOH surface roughening and AuPd coating. Leveraging their processability, PolyGraph composites are fabricated into flexible, individually isolated microneedle electrode arrays with biomimetic soft hyaluronic acid backings. These arrays demonstrate bidirectional neural interfacing capabilities, enabling both the delivery of controlled stimulation pulses in physiological buffer and high-resolution neuronal recording in murine brain slices, with machine learning-based event classification. Together, these advances establish PolyGraph as an optimal material platform for next-generation brain-computer interfaces and soft bioelectronic devices.

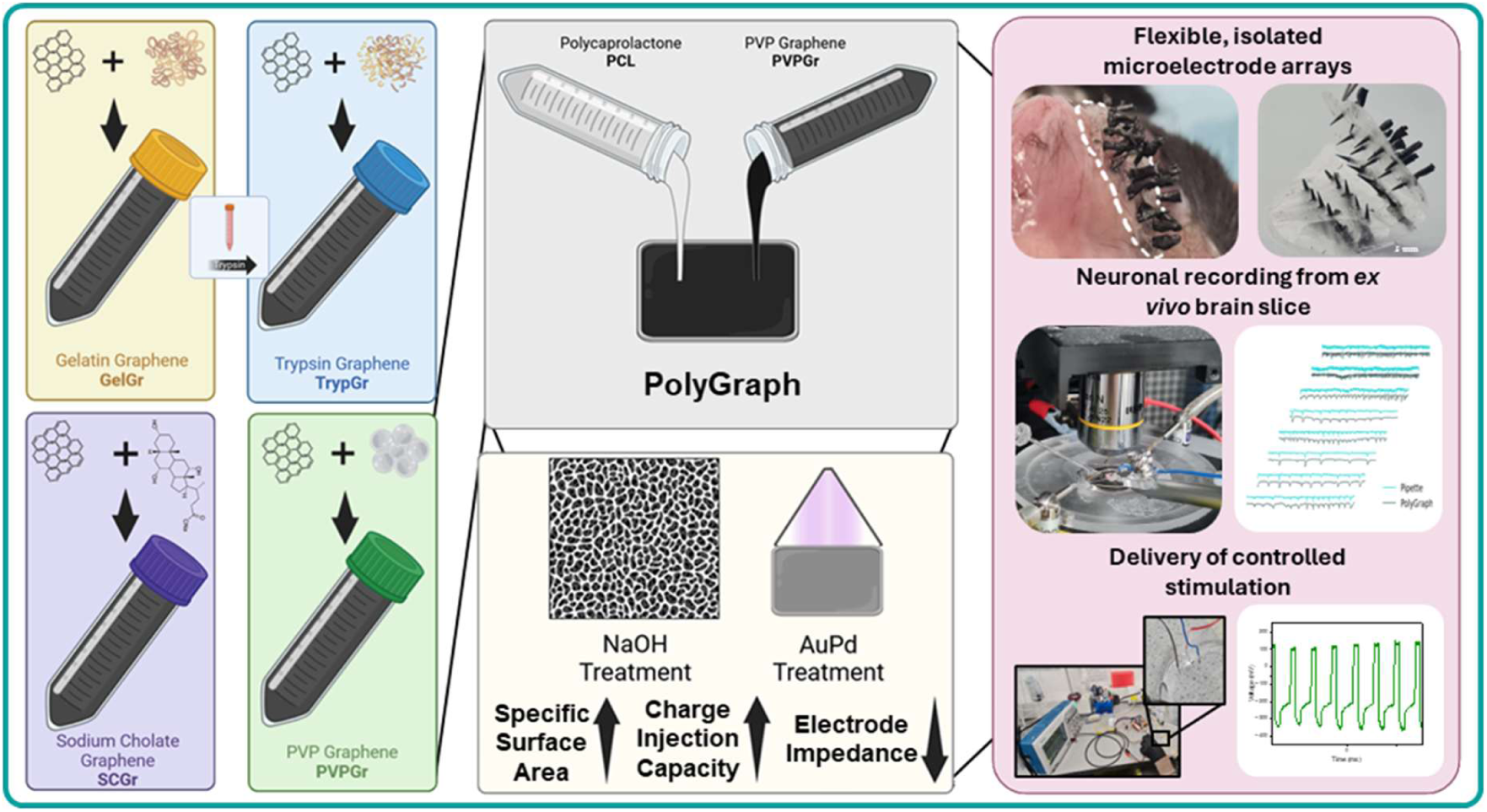

**Graphical Abstract & TOC Text** PolyGraph, a flexible graphene-polycaprolactone nanocomposite, unites conductivity, biocompatibility, and processability for next-generation neural interfaces. Fabricated into microneedle arrays with ultra-flexible backings, PolyGraph enables bidirectional neuronal recording and stimulation in brain tissue, advancing brain-computer interface (BCI) and soft bioelectronic applications.

## 1 Introduction

Over the past half-century, advances in our understanding of the central nervous system (CNS) have paralleled a revolution in neural interfacing technologies – devices that are capable of observing and modulating nerve and brain activity in real time, which have begun to revolutionise the treatment of epilepsy and Parkinson’s disease.^1–10^ Furthermore, advances in brain-computer interface (BCI) technology have the potential for profound implications in medicine and society, offering treatments for neurological disorders and paralysis, and even the prospect of seamless human-machine communication as resolution and signal interpretation improve.^2,11,12^ Although remarkable progress has been made in microneedle arrays, neural threads, and mesh electronics, these technologies remain far from joining the clinical mainstays of neural interfacing, electroencephalography (EEG)^13,14^ and functional magnetic resonance imaging (fMRI). This is despite their drawbacks, of low spatial resolution in the case of EEG,^15^ high cost and limited portability for fMRI,^16,17^ and inability to stimulate and modulate neural activity for both, leading to significant demand for high-resolution bidirectional neural interfaces. The gulf between the speed of technological development for these devices compared to their slow clinical implementation underscores the challenges involved – they must offer durable function and high resolution, without inducing scarring, infection, or device degradation.^11^ These barriers do not arise for lack of innovation on device design, signal processing, or surgical techniques, but rather due to fundamental limitations of existing neural interfacing materials.^4,7,18^

Novel electroconductive materials that combine biocompatibility, flexibility, and robust electrochemical performance are thus critical for advancing next-generation neural interfaces. Two-dimensional (2D) nanomaterials, consisting of high-aspect ratio, few-atom thick layers, are particularly well-positioned to address these challenges due to their exceptional electrical, mechanical, and morphological properties.^19–22^ As scalable, processable materials, they offer a compelling alternative to metals and doped semiconductors, which often provoke fibrotic responses, hindering widespread clinical translation.^23–25^ However, harnessing the full potential of 2D nanomaterials requires overcoming limitations of hydrophilicity and biocompatibility, typically achieved via surface functionalisation.^26–28^ Low molecular weight solvents and surfactants (e.g. N-Methyl-2-pyrrolidone, sodium cholate) offer excellent electrical properties^29–31^ but poor cytocompatibility, while biopolymers (e.g. gelatin, bovine serum albumin) improve biocompatibility and hydrophilicity, at the expense of electrical performance.^32–36^ Emerging stabilisers such as pyrene derivatives,^37,38^ polyethylene glycol, and polyvinylpyrrolidone (PVP) show promise in striking this balance, though performance remains highly application-dependent.^39–43^

Pairing 2D materials with biocompatible polymers offers a promising route to achieving electrical performance, bioactivity, and durable mechanical properties simultaneously^44–46^. Polycaprolactone (PCL), a biocompatible and processable thermoplastic,^47^ is one such polymer, enabling 3D printing, casting, and molding of nanocomposites.^48^ However, few studies have explored the use of nanomaterial composites for neural interfacing electrodes when compared to both traditional and purely 2D systems,^19,49^ despite their potential to address many of the issues present in existing devices. In this work, we develop PolyGraph, a soft neural interface material based on a novel PVP-stabilised graphene formulation, embedded within a PCL matrix. This composite exhibits exceptional electrochemical performance, robust biocompatibility, and mechanical flexibility – key attributes that make it well-suited for minimally invasive bidirectional neural electrodes.

Nonetheless, a nanocomposite, however advanced, is far from a functional device. Effective neural interfacing requires integration of novel materials into microelectrode,^50,51^ microneedle,^52,53^ and soft biohybrid^54,55^ architectures, which have dramatically improved both *in vitro* and clinical neural interfacing capabilities. The smaller addressable volume of these micro-scale electrodes enables more precise control of external devices (e.g. prosthetics, communicators, computers), more accurate interpretation of neural signals, and a deeper understanding of CNS function.^56,57^ In this study, we fabricate a PolyGraph microneedle-based electrode array with PDMS sheathing to reduce electrode size, and a flexible, biomimetic, immunomodulatory hyaluronic acid backing for conformability with neural tissues, decoupling the micromotion of the brain from the device.^23,58^ We demonstrate its capacity for bidirectional neural interfacing, recording neuronal activity from murine brain slices, and delivering stimulation pulses in physiological conditions.

This study introduces PolyGraph, a soft, flexible, biocompatible, and electrochemically optimised graphene-polymer composite developed to address the trade-offs of conductivity, biocompatibility, and mechanical mismatch in neural interfaces. We first identified polyvinylpyrrolidone (PVP) as an optimal stabiliser for liquid-phase exfoliation of high-quality graphene, balancing conductivity and cytocompatibility. This PVP-stabilised graphene was incorporated into PCL to create versatile, highly conductive nanocomposites, which were subsequently engineered for enhanced electrochemical performance via NaOH surface roughening and AuPd coating. Leveraging the processability of these composites, we fabricated a multi-channel microneedle-based electrode array with a soft, biocompatible hyaluronic acid backing for conformability with neural tissues. Finally, we demonstrated the bidirectional functionality of PolyGraph through extracellular neuronal recording in *ex vivo* murine brain slices, coupled with machine learning-based classification of neural events to enable real-time interpretation, and stimulation in physiological medium. By simultaneously overcoming key limitations of existing microelectrode designs and uniting softness, biocompatibility, and versatile, scalable fabrication, PolyGraph establishes itself as a promising platform for next-generation brain-computer interfaces and soft bioelectronics, such as neural meshes, threads, microelectrode arrays, and deep brain stimulation devices.^59–67^

## 2 Results and Discussion

### 2.1 Optimisation of graphene formulation from range of candidate exfoliations

#### 2.1.1 Physical Characteristics

For bioelectronics applications, there is a strong clinical need for soft materials with high conductivity and excellent biocompatibility. We first aimed to overcome the trade-off between these two properties by identifying an optimal graphene stabiliser for use in the CNS. Complicating this optimisation is the fact that many biocompatible stabilisers used to reduce nanomaterial cytotoxicity also significantly hamper their conductivity, due to steric hindrance and the disruption of nanosheet junctions.^68,69^ To address this, we evaluated four stabilisers for graphene liquid-phase exfoliation (LPE) (**Figure S1**A).

Optimization began with gelatin-coated graphene (GelGr), which exhibited low conductivity (12 S/m), likely due to junction disruption by gelatin (**Figure 1**A), with composites fabricated using this graphene (**Figure S2**A) failing to reach the range required for low-impedance, high-resolution neural interfacing.^70^ To probe the effect of stabiliser molecular weight, we digested the gelatin layer enzymatically using trypsin^71^ (TrypGr). Conductivity measurements revealed a significant improvement (Figure 1B), supporting our hypothesis that thinner stabiliser coatings improved nanosheet junction quality and network conductivity. Thermogravimetric (TGA) analysis corroborated this, showing reduced stabiliser content in TrypGr (0.82%) compared to GelGr (4.64%)(**Figure S5**), while scanning electron microscopy (SEM), revealed disruption of the gelatin coating (inset, Figure 1A). Based on this evidence linking stabiliser molecular weight to conductivity, we next tested two well-established smaller-molecule exfoliants: sodium cholate (SC)^72^ and polyvinylpyrrolidone (PVP).^40,43^ Both yielded significantly higher conductivities and thinner, higher quality nanosheets than GelGr and TrypGr (Figure 1A, 1B).

**Figure 1.**
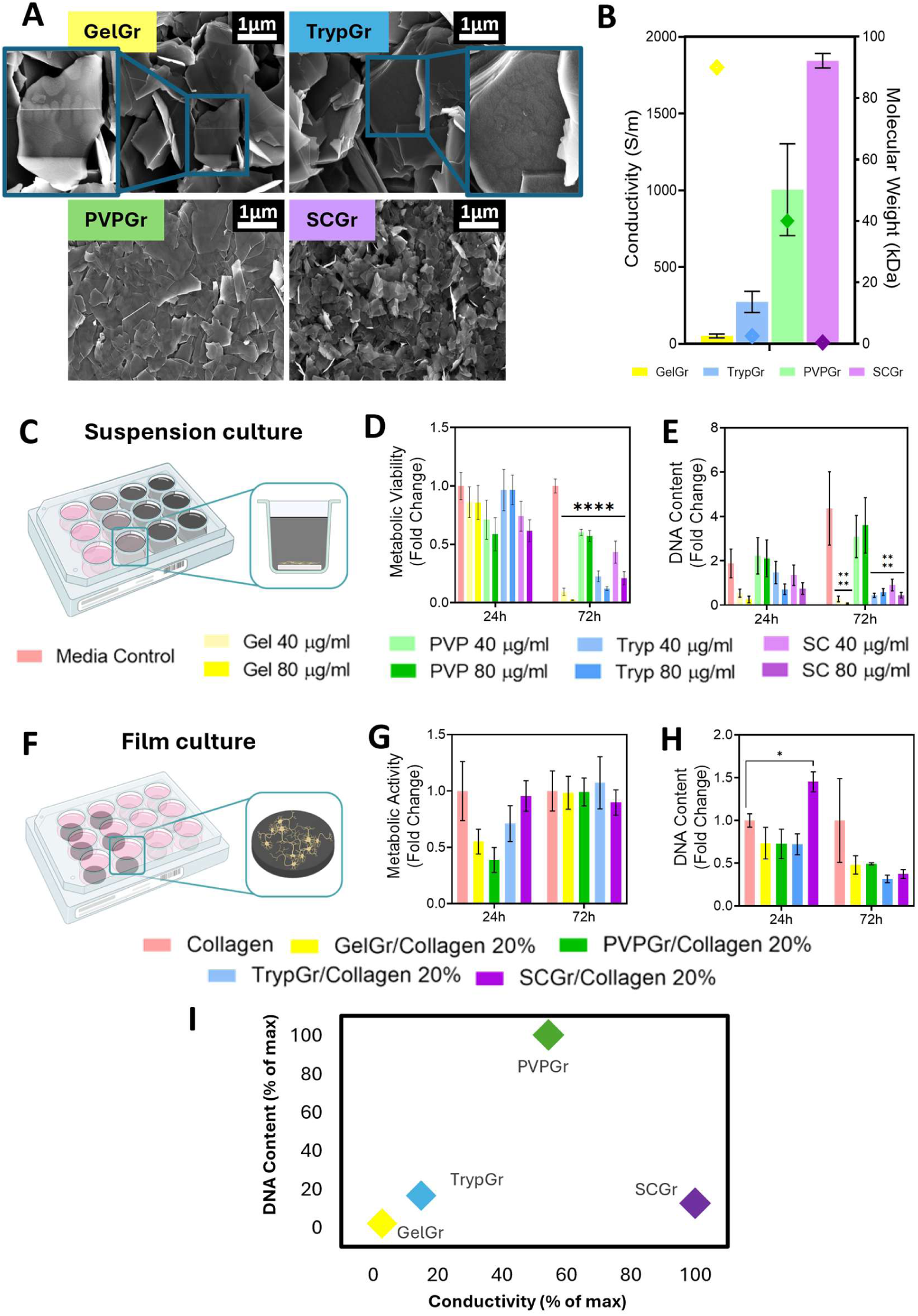
Physical characterisation of graphene candidates: **A)** SEM imaging of GelGr, TrypGr, PVPGr and SCGr formulations. Inset highlights disruption of gelatin layer by trypsin treatment. Scale bars 1 μm. **B)** Conductivity of graphene films (bars), with molecular weights of stabilisers indicated by diamonds. **C)** Schematic of suspension biocompatibility assays. **D)** Metabolic activity and **E)** DNA content of NSC-34 cells grown in suspension with 40 or 80 μg/mL of each graphene formulation. **F)** Schematic of collagen film biocompatibility assays. **G)** Metabolic activity and **H)** DNA content of NSC-34 cells grown on collagen films containing 20 wt% graphene. **I)** Plot of cellular survival versus conductivity, expressed as percentage of maximum DNA content and conductivity for each graphene type. Significances: *p < 0.05, **p < 0.01, ***p < 0.001, ****p < 0.0001.

#### 2.1.2 Biological Characteristics

Having compared their physical characteristics, we next assessed candidate biocompatibility. To robustly examine the relative cytotoxicity of the nanoparticles, mouse motor neurons (NSC-34s) were cultured with high-concentration suspensions of each material (80 μg/mL) for 3 days (Figure 1C). This approach models an acute release of nanomaterial into the body – the highest risk scenario for cellular internalisation, wrapping, and other damaging cell-nanomaterial interactions.^73^ Cellular health and survival, assessed by metabolic activity (Figure 1D) and DNA content (Figure 1E), revealed significant cytotoxicity for all formulations, with the exception of PVPGr, which maintained healthy neuronal growth and metabolic activity.

This striking result underscores the critical impact of stabiliser choice on nanomaterial biocompatibility. PVP, a commonly used biopolymer,^39,40^ likely enhances biocompatibility due to steric shielding, protecting cells from direct contact with the sharp nanomaterial, and prevention of protein adsorption.^74,75^ This is consistent with findings for aromatic group-based stabilisers such as bis**-**pyrene^26,37^ and polyethylene glycol (PEG).^74,76,77^

Recognising that immobilization of nanomaterials within a polymer matrix to form a nanocomposite can significantly enhance biocompatibility, we next assessed neuronal viability on collagen**-**graphene composite films with each candidate graphene formulation (20 wt%, Figure 1F). In this case, the collagen matrix and reduced concentration of free graphene led to no significant reduction in cell population after 72 h (Figure 1G, S1B), suggesting that the polymer matrix effectively mitigated cell-nanomaterial interactions, reducing cytotoxic effects. The data thus far highlights PVP-stabilised graphene (PVPGr) as the optimal candidate for further study, exhibiting a strong balance of biocompatibility with supra-physiological conductivity (Figure 1I), positioning it as a promising material for use in the CNS.

### 2.2 PVP-stabilised graphene (PVPGr) characterisation

To confirm the robustness of the properties of PVP-stabilised graphene (PVPGr) across a range of nanosheet morphologies and exfoliation parameters, we exposed mouse motor neurons to PVPGr formulations derived from distinct LPE protocols (**Methods Table 1**). All formulations maintained largely stable neuronal metabolic activity (**Figure 2**A) and DNA content (Figure 2B), outperforming sodium cholate-stabilised graphene (a commonly used stabilising surfactant). This finding underscores the broad biocompatibility of PVPGr, and its relative independence from the precise exfoliation protocol.

**Figure 2.**
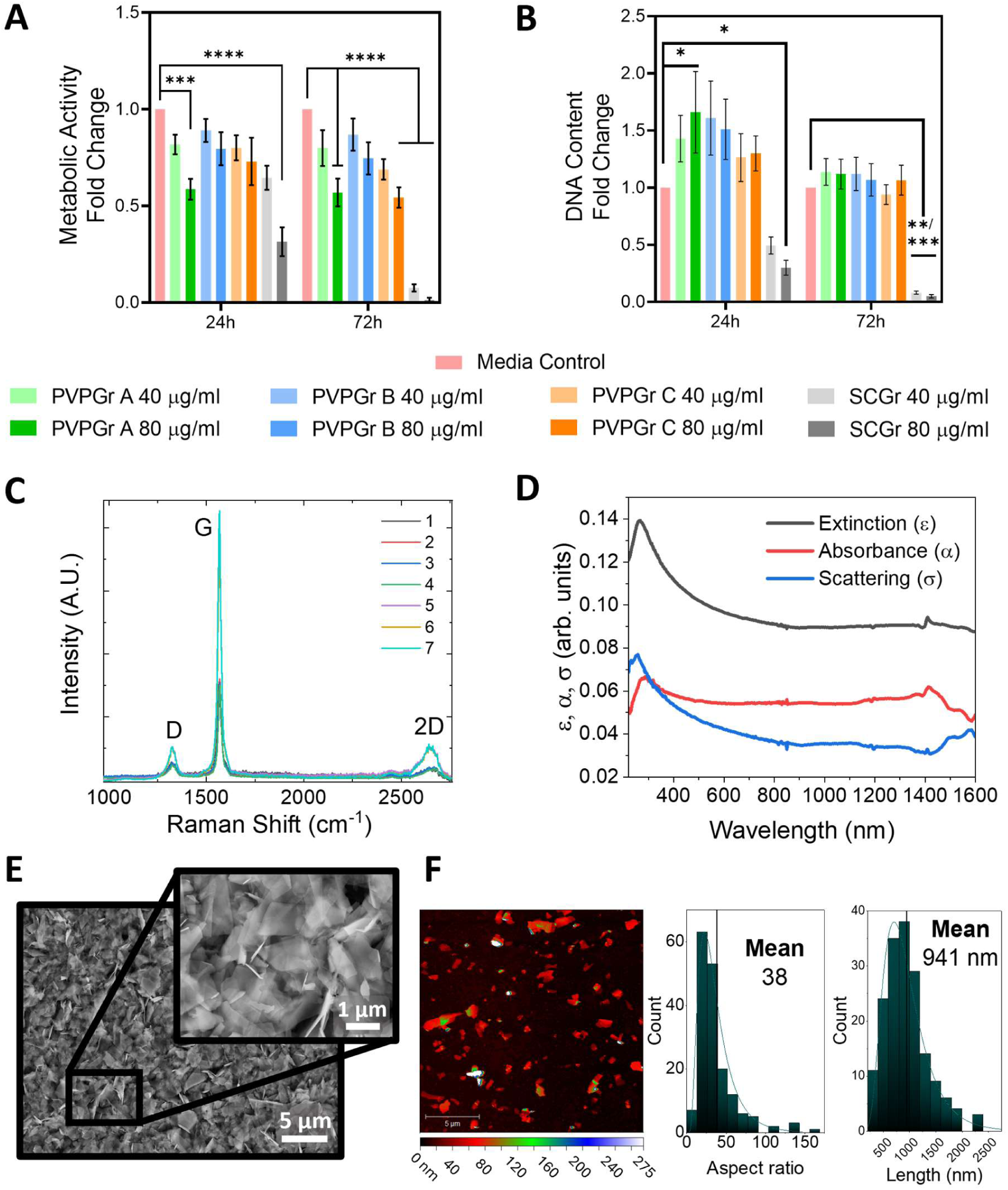
Physical & biological characterisation of PVP-stabilised graphene (PVPGr): A-B) Metabolic activity (A) and DNA content (B) of NSC-34 cells in suspension with PVPGr formulations, showing robust biocompatibility. **C)** Raman spectra of PVPGr, with D, G, and 2D peaks characteristic of minimally-defected graphene sheets. **D)** UV-Vis spectrum of PVPGr dispersion, with absorption and scattering features typical of graphene and nanomaterials. **E)** SEM images of a PVPGr network, with inset showing thin, overlapping nanosheets. Main scale bar 5 μm, inset scale bar 1 μm. **F)** AFM analysis of PVPGr, showing high aspect ratio and lateral size. Scale bar 5 μm. Significances: *p < 0.05, **p < 0.01, ***p < 0.001, ****p < 0.0001.

To assess the basal plane quality of PVPGr, Raman spectroscopy was next performed (Figure 2C). Plotting the intensity ratio of the D and G peaks 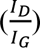 against the full-width-half-maximum of the G peak (FWHM_G_), revealed no correlation (**Figure S4**C) strongly suggesting that the observed D peak and associated disorder are not caused by basal plane imperfections, but rather by edge defects,^78^ consistent with previously reported LPE graphene inks.^79,80^ Building on this, we can estimate the average lateral size of the nanosheets (<L>) using the relationship 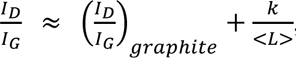, with *k* = 0.17.^81^ Using 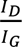 ≈ *0.05 (± 0.05)* for our graphite^82^ and an average 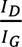 of 0.15 for the exfoliated graphene flakes, we estimated the lateral size of the nanosheets as 1.13 – 3.4 µm.

UV-Vis spectroscopy showed the characteristic spectra of few-layer graphene,^83,84^ with a pronounced absorbance peak occurring at 284 nm (Figure 2D), alongside a significant degree of scattering, typical of nanomaterials.^85^ SEM imaging revealed thin, evenly sized nanosheets (Figure 2E), while atomic force microscopy (AFM) analysis yielded a mean nanosheet length of 0.941 µm and aspect ratio of ∼38 (**Figure** 2F, **S5**), aligning with Raman estimates. Collectively, these results confirm the production of high-quality, biocompatible graphene sheets ideally suited for soft bioelectronics applications.

### 2.3 Manufacture and optimisation of PCL-graphene composite (PolyGraph) materials

Polymer-free nanomaterial networks often suffer from fragility, as their cohesion arises due to weak van der Waals interactions, leaving them prone to delamination and degradation.^86^ To overcome this, we developed a composite system harnessing the mechanical robustness and processability of polymers, while preserving the exceptional properties of graphene. The chosen polymer matrix needed to meet several criteria, including low molecular weight to minimize steric hindrance and maintain nanosheet conductivity, along with biological stability and biocompatibility. Polycaprolactone (PCL), a biocompatible thermoplastic widely used in biomedical applications,^47,87^ was selected for its versatile processability (3D printing, casting, and molding).^48^ Its relatively low modulus (∼350 MPa^88^), two orders of magnitude lower than gold^89^ and three orders of magnitude lower than platinum or silicon,^90,91^ is expected to enhance cellular adhesion and proliferation, while reducing stiffness-induced foreign body response.^92^

The optimized manufacturing process (Figure 3A) produced flexible PCL-graphene (PolyGraph) composite films with minimal aggregation. To balance conductivity and mechanical properties, we assessed the percolation threshold in a graphene loading range of 0.125 vol% to 20 vol% (Figure 3C). Percolation was reached at ∼1.45 vol%, with a critical exponent of 3.1, higher than the value of *t = 2* predicted by theory^93,94^ – this may be explained by the broadening of the distribution of both nanosheet and junction resistances due to variability in polymer coating and sheet quality (Figure S2).^93–96^ Based on these results, 10 vol% (PolyGraph10%, ∼36 S/m) and 20 vol% (PolyGraph20%, ∼228 S/m) were chosen for further study, as they exhibited the best balance between electrical and mechanical properties, suitable for CNS applications.

**Figure 3.**
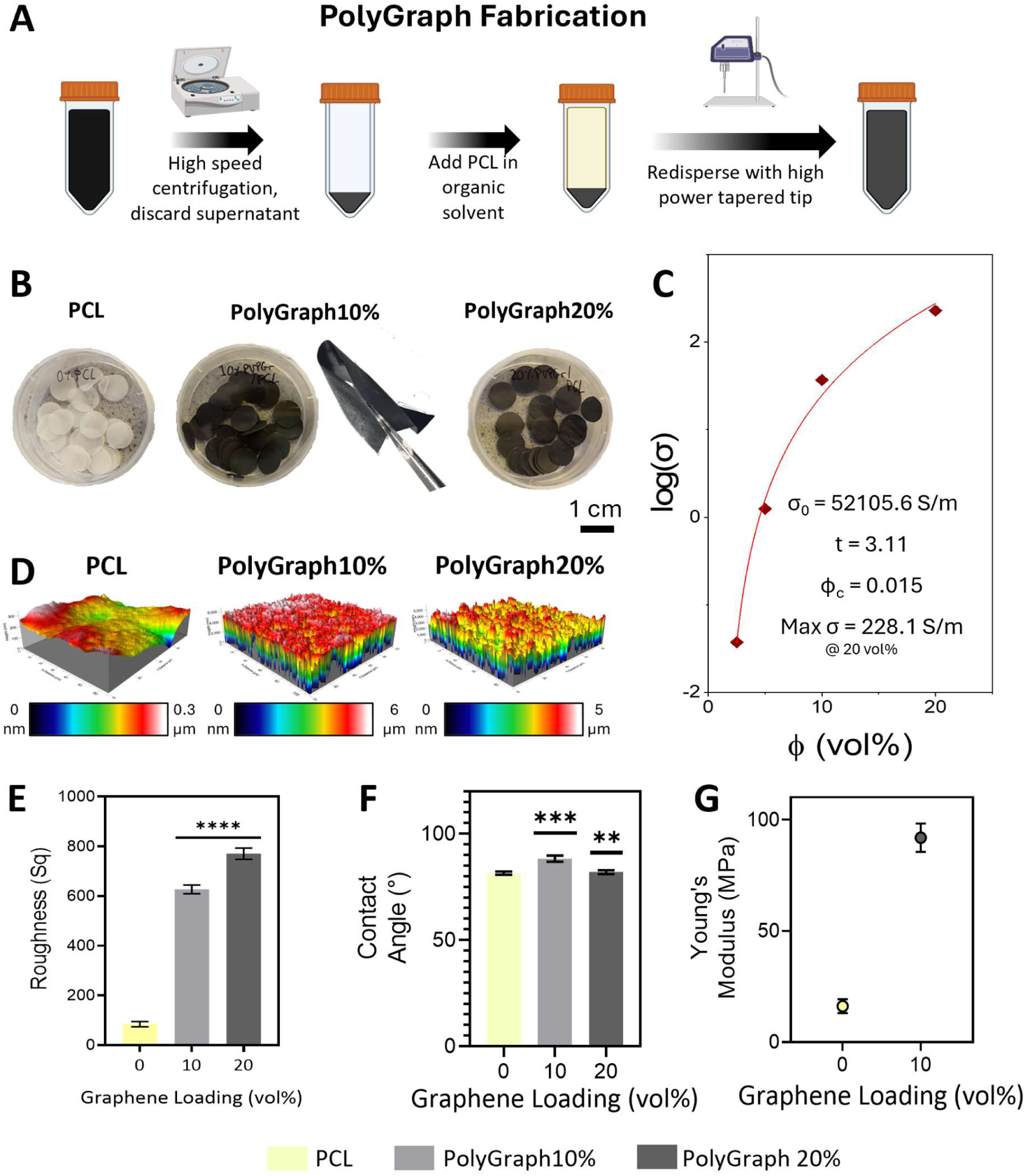
Physical characterisation of PolyGraph composites: **A)** Schematic of PolyGraph fabrication process. **B)** Optical images of PCL and PolyGraph. **C)** Percolation curve for PolyGraph conductivity with fitting parameters (percolation threshold Φ_c_, network conductivity σ_0_, critical exponent t). **D-E)** White light interferometry data (D) for PCL and PolyGraph, indicating increased roughness with graphene loading (E). **F)** Contact angle measurements for PolyGraph composites, indicating a slight but significant increase in hydrophobicity with graphene loading. **G)** Young’s modulus of PolyGraph, showing increased tensile strength with graphene loading. Significances: *p < 0.05, **p < 0.01, ***p < 0.001, ****p < 0.0001.

Both formulations yielded homogeneous, flexible films (Figure 3B). Surface roughness increased with graphene loading (Figure 3D, 3E), disrupting the smooth polymer surface in a manner known to promote cellular adhesion^97–99^ and enhance electrochemical performance.^100,101^

Mechanical testing showed increased Young’s modulus with graphene addition (Figure 3G), accompanied by reduced ductility **(Figure S6)**. This decrease likely arises due to the Rule of Mixtures, where weak inter-flake interactions dominate at high loadings, reducing composite strength and ductility, while also changing its viscoelastic properties.^102^ Contact angle measurements (Figure 3F) revealed a slight decrease in hydrophilicity for PolyGraph samples. Finally, TGA confirmed graphene loadings matched target fractions (**Figure S4**A, S4B). Collectively, these findings position PolyGraph as a flexible, mechanically robust, and electrically optimised platform for advanced soft bioelectronics.

### 2.4 Response of neuronal and glial cells to PolyGraph composites

#### 2.4.1 Neuronal response

Some of the most critical factors in electrode design for biomedical applications are cytocompatibility and immune response, as they dictate the safety and lifetime of implanted devices.^103^ We therefore investigated the ability of neuronal cells to grow on PolyGraph without inducing adverse responses. First, the ability of human neuroblastoma-derived SH-SY5Y cells to grow and colonise PolyGraph substrates was assessed. Neurons grown on all tested films demonstrated increases in metabolic activity and DNA content after 3 days of growth, indicative of cell proliferation and surface colonisation, while showing no significant changes in metabolic activity (**Figure 4**A) or DNA content (Figure 4B) compared to control substrates. Next, to evaluate biocompatibility with a more physiologically relevant cell type, human induced pluripotent stem cell (iPSC)-derived neurons were seeded on the PolyGraph films and allowed to grow for 14 days. PolyGraph10% supported the extension of neurites as well as significantly increased cellular coverage of the films (**Figure** 4C, 4D, **S7**), potentially due to enhanced surface roughness and substrate conductivity.^98^ PolyGraph10% also showed trends towards increased metabolic activity and maximum neurite length (**Figure S8**A, S8B). Interestingly, PolyGraph20% reduced neuronal populations (Figure 4D), suggesting a critical graphene loading exists beyond which adverse effects may arise due to increased exposure to nanosheet edges (**Figure 5**A) and resulting mechanical stress on cells.^104^ Mouse primary cortical neurons cultured on PolyGraph confirmed its broad biocompatibility, even at higher loadings (**Figure S9**). These findings emphasize the importance of optimizing the graphene loading, as it is difficult to predict the biological behaviour of nanomaterials in different configurations due to complex physical and cellular interactions.

**Figure 4.**
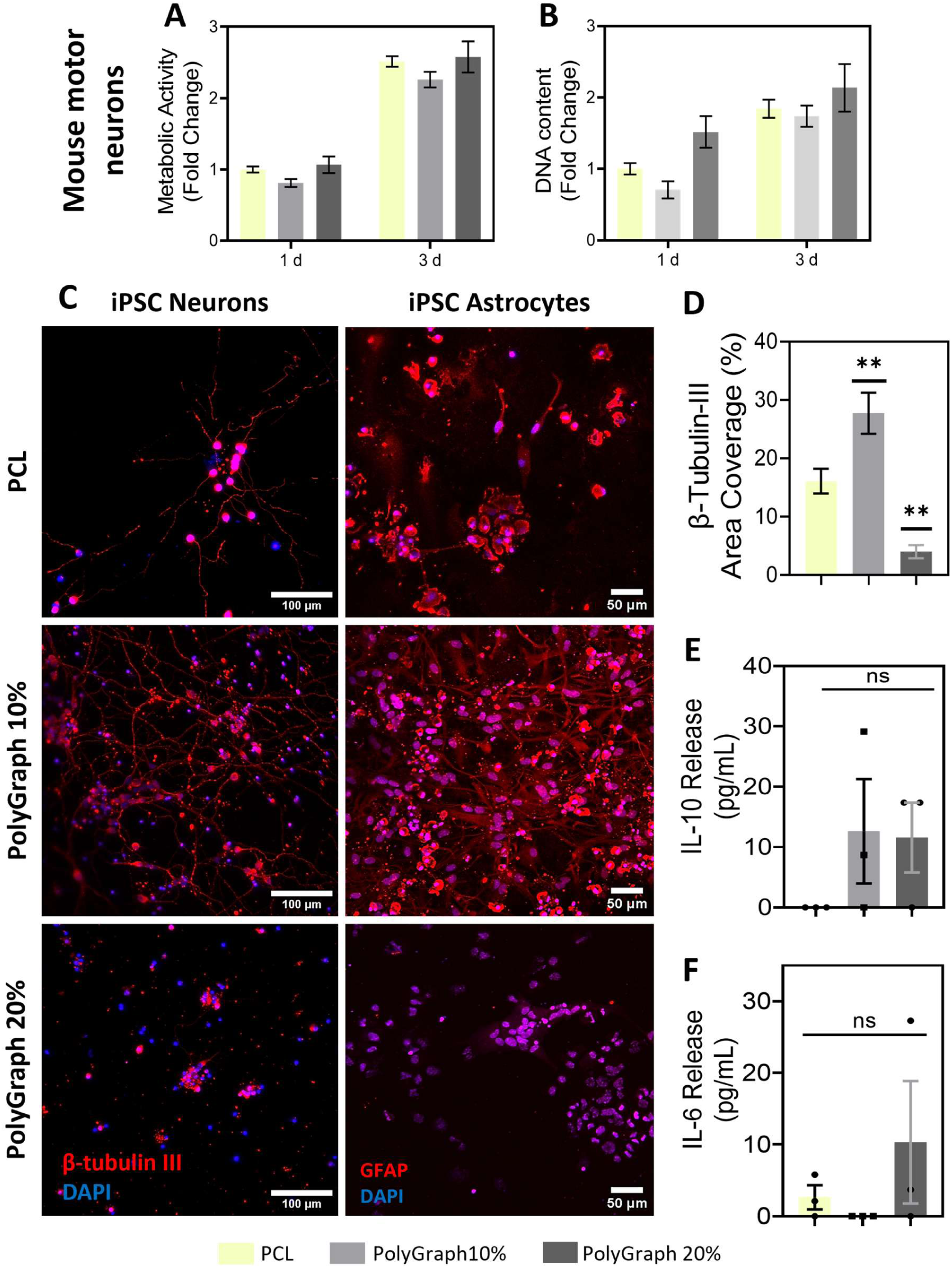
Biological & immunological characterisation of PolyGraph: A-B) Metabolic activity (A) and DNA content (B) of SH-SY5Y neurons cultured on PolyGraph over 3 days. **C)** Representative immunofluorescence images of iPSC-derived neurons and astrocytes after 14 days on PolyGraph, demonstrating robust survival, healthy morphologies, and extensive neurite outgrowth across the surface of PolyGraph10%. Scale bars: neurons 100 μm, astrocytes 50 μm. **D)** Quantification of β-III tubulin coverage, indicating increased neuronal coverage on PolyGraph10%. **E-F)** Cytokine release from iPSC-derived astrocytes: E) anti-inflammatory IL-10, and F), pro-inflammatory IL-6. Significances: *p < 0.05, **p < 0.01, ***p < 0.001, ****p < 0.0001.

**Figure 5.**
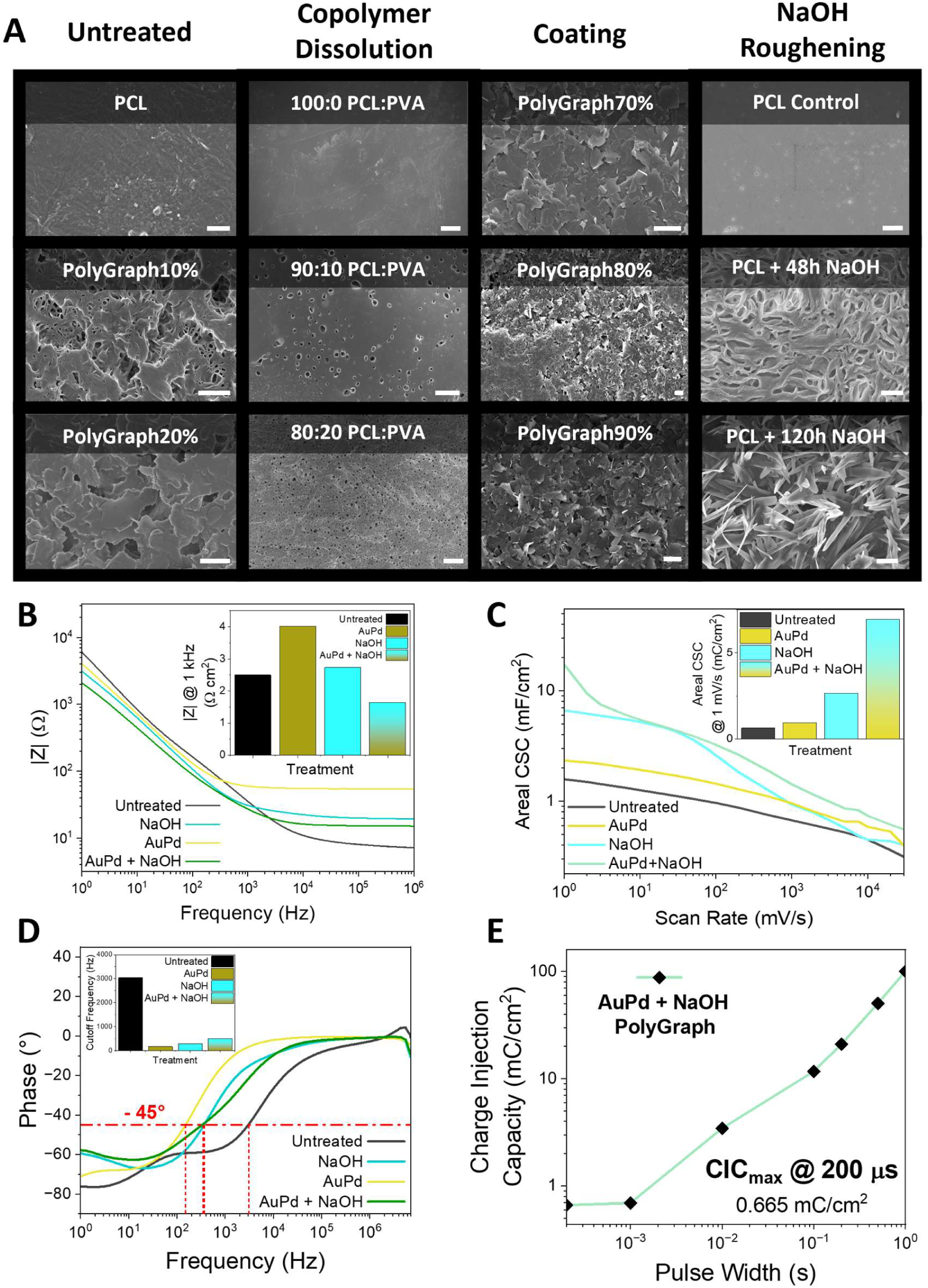
Electrical optimization & characterisation of PolyGraph: **A)** SEM images of PCL and PolyGraph following application of roughening treatments, from left to right: untreated PolyGraph, copolymer dissolution with PVA, spray coating with high-loading PolyGraph, NaOH roughening. Scale bars 2 μm, except for copolymer dissolution, which are 100 μm. **B)** Electrochemical impedance spectroscopy, with Bode plots showing reduced impedance after AuPd coating and NaOH treatment of PolyGraph10%. Inset: areal impedance at 1 kHz. **C)** Cyclic voltammetry showing significantly increased charge storage capacity following NaOH and AuPd treatment. Inset: CSC at 1 mV/s scan rate. **D)** Cut-off frequency analysis derived from Bode phase plots, with significant decrease observed following surface treatment. **E)** Voltage transients analysis determining charge injection capacity (CIC), with surface-treated PolyGraph10% achieving a CIC of 0.665 mC/cm^2^ for a 200 μs pulse.

#### 2.4.2 Glial response

The foreign body response, driven by reactive responses of CNS-resident astroglial cells, is another critical determinant of long-term electrode performance, as even low levels of fibrotic scarring can severely limit device lifetime and function.^105^ We assessed the response of naïve iPSC-derived astrocytes to PolyGraph, finding that PolyGraph10% supported healthy stellate morphologies with long processes (Figure 4C).^106^ ELISA revealed no significant changes in the release of anti-inflammatory IL-10 or pro-inflammatory IL-6 from cells grown on PolyGraph compared to collagen, indicating an absence of cell polarisation (Figure 4E & 4F).

A slight trend towards reduced LDH release on PolyGraph films was observed at day 7, with no impact on the metabolic activity of the astrocytes (Figure S8D, S8C). Together, these data indicate that PolyGraph10% is capable of supporting astroglial growth without inducing damaging foreign body responses from physiologically relevant cells. Combined with its mechanical, electrical, and biological advantages, these findings position PolyGraph10% as a flexible, biocompatible platform for long-term neural interfaces, and justify its selection for further device development.

### 2.5 Electrochemical characterisation of PolyGraph

With the biocompatibility of PolyGraph10% with key neuronal and glial cell types established, focus was shifted to its electrochemical performance, to assess its suitability for neural stimulation and recording. Key parameters – surface impedance (*Z*), charge storage capacity (CSC), cut-off frequency (*f_cut-off_*) and charge injection capacity (CIC) – were assessed in untreated films, and following surface modification. These modifications were employed to enhance electrochemical performance by increasing the electrochemical surface area (ESA)^100,101^ and decreasing the impedance (Figure 5A). These parameters impact the signal-to-noise ratio (SNR), efficiency of stimulation, resolution of recording, and other key outputs,^103^ and can be tuned by material choice, post**-**treatment, and device morphology.^107^

Untreated PolyGraph10% exhibited a surface impedance of 2.5 Ω cm^2^ at 1 kHz (Figure 5B) and a charge storage capacity (CSC) of 1.6 mC/cm^2^ at a scan rate of 1 mV/s (Figure 5C), with a cutoff frequency of 3,040 Hz. The electrochemical potential window, assessed by chronoamperometry, was ±0.5 V (Figure S10B), narrower than the thermodynamic water window (∼±0.6 V).^108^ These values, while within literature ranges for impedance and CSC (0.01-1 Ω cm^2^ & 0.2-100 mC/cm^2^ ^109–111^), and sufficient for large electrodes and stimulatory applications, fall short of the requirements for high-resolution neural recording.^18,112^

Several post-treatments were tested (Figure 5A). Channel formation in PCL using polyvinyl alcohol (PVA) as a sacrificial copolymer increased porosity, but proved inconsistent. Spray coating with high-loading PolyGraph did not significantly alter CSC (**Figure S11**A), and the reduction in polymer binding led to a risk of electrode delamination. Both approaches were therefore excluded from further development.

Sodium hydroxide (NaOH) treatment, a well-established technique for roughening PCL,^113–115^ dramatically improved electrode performance. CSC increased 11-fold to 17.3 mC/cm^2^ (Figure 5C), corresponding to a volumetric capacitance of 787.4 mF/cm^3^ (Figure S11B). SEM imaging confirmed the roughened surface, and literature suggests such surface modification may also enhance cellular adhesion and survival.^97,99,113,115^ To further reduce impedance, a thin

AuPd coating was applied.^116^ This treatment decreased the impedance of the composite materials significantly when used in conjunction with NaOH treatment, from 2.5 Ω cm^2^ to 1.6 Ω cm^2^ (Figure 5B), evidence of the beneficial synergistic effect of these treatments. The safe electrochemical potential window also widened to ±0.7 V (Figure S10C), supporting higher current delivery without electrolysis. Cut-off frequency was reduced to 495 Hz (Figure 5D), ideal for action potential recording,^107^ and CIC measured using a three-electrode setup reached ∼100 mC/cm^2^ for 1 s pulses, with ∼0.65 mC/cm^2^ at 200 μs (Figure 5E, S12B). These values compare favourably to literature reports of CIC in neural electrodes (0.1 – 80 mC/cm^2^).^110,117–121^ Limitations in accounting for pulse width and access voltage suggest these are conservative estimates.^107^

These findings demonstrate that NaOH and AuPd treatments synergistically enhance the electrochemical performance of PolyGraph10% electrodes, enabling safe and efficient neural stimulation and high-resolution recording. This opens new pathways for developing minimally invasive neural devices using electrochemically optimised PolyGraph, capable of addressing the complex demands of clinical and research applications. Building on this, materials with proven biocompatibility and electrochemical performance hold significant potential across bioelectronics, including in small site-size neural interfaces, as explored in the next sections.

### 2.6 Microneedle-based Neural Interface Fabrication

Having successfully developed and optimised PolyGraph10% composites with enhanced electrochemical properties via NaOH roughening and AuPd coating, as a proof of concept, we next utilised these PolyGraph10% composites to fabricate a flexible microneedle-based neural interface. Effective high-resolution neural interfacing requires miniaturization, flexibility and high channel count,^18^ typically achieved using expensive fabrication techniques such as photolithography and chemical vapour deposition.^122^ Here, we demonstrate a scalable, low-cost mold-based fabrication protocol (**Figure 6**A), yielding flexible microneedle arrays with individually isolated electrodes. These electrodes are connected by a hyaluronic acid extracellular matrix (ECM) backing that mimics the primary matrix component of the surrounding CNS,^23,58^ improving integration with the brain surface and decoupling implant micromotion from that of the brain (Figure 6B).

**Figure 6.**
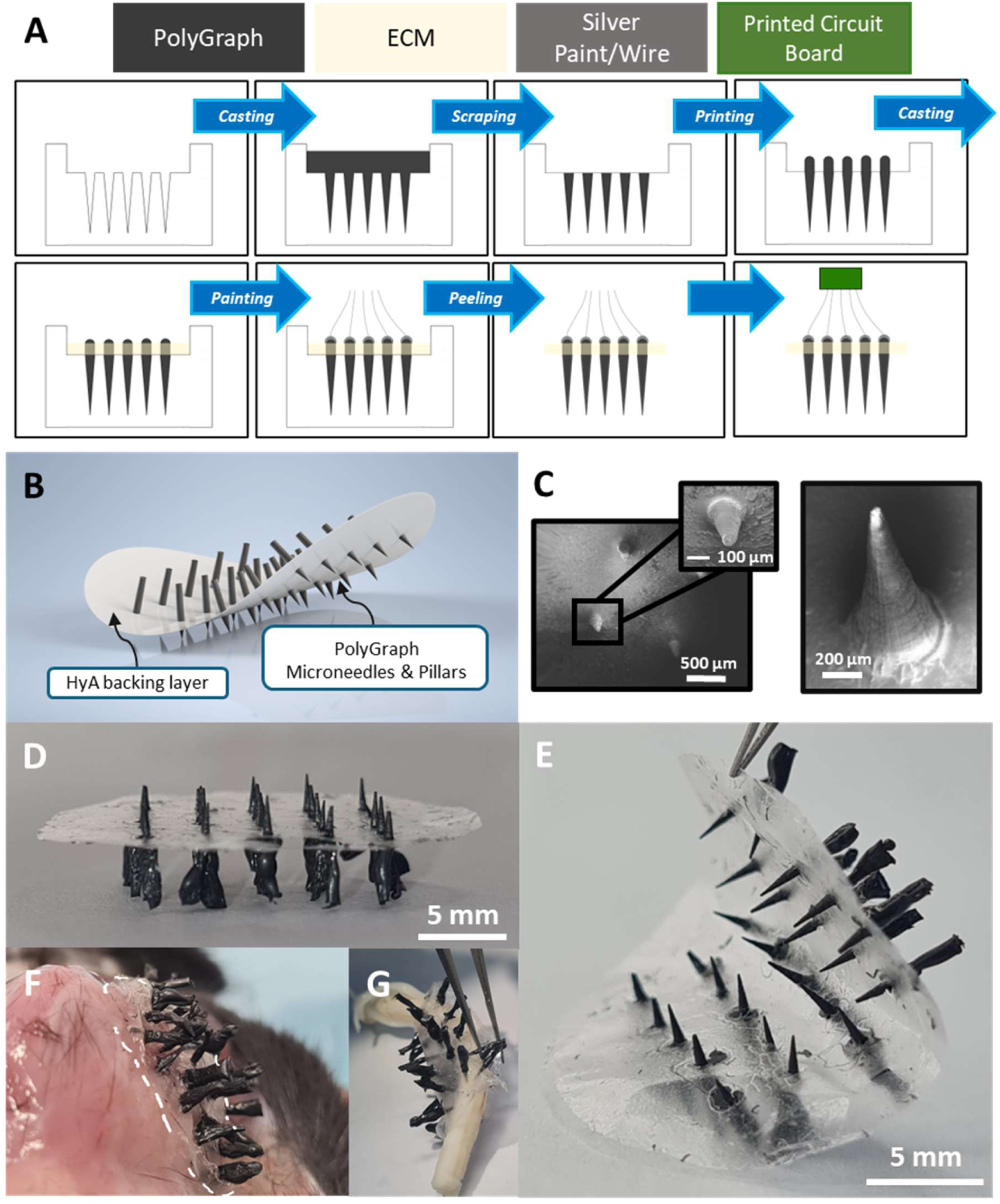
Fabrication of flexible, electrically isolated microneedle arrays: **A)** Schematic of PolyGraph microneedle array fabrication workflow, leverage composite processability for dry casting and 3D printing. **B)** Render of final device, showing PolyGraph microneedles and pillars anchored within a flexible hyaluronic acid (HyA) backing layer. **C)** SEM images of individual PolyGraph microneedles at various magnifications, highlighting sharp tips and uniform geometry. **D-G)** Optical images of the fabricated flexible, isolated microelectrode array: D), isolated array, E) demonstrating mechanical flexibility of backing, F) penetrating mouse dorsal tissue (white outline indicates hyaluronic acid backing), and G) wrapping around a rat spinal cord. Scale bars as shown.

PolyGraph10% microneedle arrays were melt-cast using custom silicone mould**s**, then combined with 3D-printed PolyGraph10% to produce isolated needle-pillar electrode pairs (**Figure S13**A, S13B). SEM (Figure 6C) and white light interferometry (WLI, Figure S13C) confirmed high-quality microneedles with ∼50 µm tips. The 3D printability of PolyGraph10% also enabled custom scaffold and bioelectronic circuit fabrication (Figure S13D-F).

To minimise mechanical mismatch and prevent fibrotic encapsulation, the microneedles were linked via a thin, flexible hyaluronic acid backing. This freestanding, highly flexible microelectrode array (Figure 6D, 6E) is capable of conforming to complex tissue surfaces including mouse muscle (**Figure** 6F, **S14**B) and rat spinal cord (Figure 6G, S14A, S14C), unlike traditional rigid electrodes^103,123^.

Mechanical testing confirmed the backing’s softness, with forces remaining below the 0.01 N detection limit of our tensile tester, approaching a modulus comparable to native CNS tissue (<1 kPa).^23^ This softness is critical for reducing micromotion-induced chronic inflammatory responses upon implantation.

These advances establish PolyGraph10%-based microneedle arrays as a scalable, flexible, and biocompatible platform for minimally invasive, high-resolution neural interfaces, capable of addressing both central and peripheral targets.

### 2.7 Bidirectional recording and stimulation in *ex vivo* brain tissue using PolyGraph10% electrodes

Finally, to demonstrate the potential for this PolyGraph10% device as a functional bidirectional neural interface, we assessed its ability to deliver stimulation regimes in physiological buffer, and to record neuronal activity in *ex vivo* acute murine brain slices. Individual microneedles, fabricated as described above and insulated with PDMS to expose only the tip^18,112^ (**Figure 7**A, **S15**), were integrated into an electrophysiological recording system. Compressive testing confirmed that the microneedles could withstand typical microneedle insertion forces into brain tissue (0.7 – 40 mN),^124^ with a maximum compressive force of 0.36 N sustained before buckling (**Figure S13**G).

**Figure 7.**
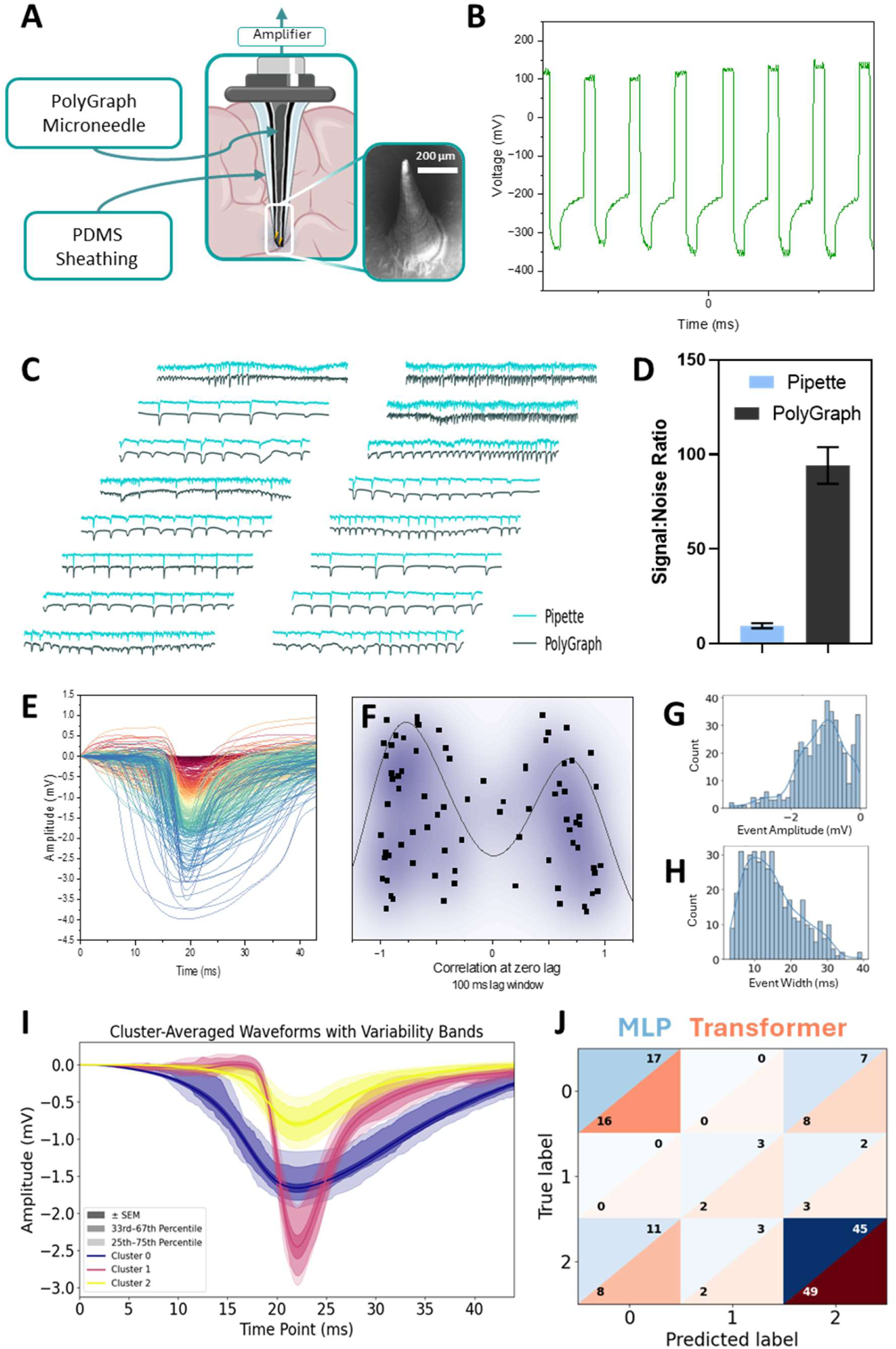
Electrophysiological recording using PolyGraph microelectrodes: **A)** Schematic of a PolyGraph microneedle electrode sheathed in PDMS to reduce active electrode site size. Inset: SEM image of sharp microneedle tip. **B)** Biphasic stimulation waveform delivered using PolyGraph microneedle via IonOptix pulse generator and recorded via oscilloscope. **C)** Neuronal activity traces from simultaneous recordings with a standard pipette electrode (blue) and a PolyGraph microelectrode (black), showing correlated neuronal activity. **D)** Signal-to-noise ratio (SNR) of recorded traces, with PolyGraph electrodes yielding SNR suitable for high-resolution recording. **E)** Overlay of aligned neuronal events recorded by PolyGraph, displaying typical local field activity event morphology. **F)** Correlation analysis between events recorded by pipette and PolyGraph electrodes, showing strong agreement. **G-H)** Event feature distributions for PolyGraph recordings: amplitude (G) and width (H). **I)** Cluster-averaged waveforms with variability bands from k-means clustering of PolyGraph event traces, revealing distinct neuronal event types. **J)** Confusion matrix for MLP neural network and simple transformer-based classifiers, showing high classification accuracy of events into k-means derived clusters.

First, PolyGraph microneedles successfully delivered biphasic stimulation pulses in phosphate-buffered saline, demonstrating charge injection capability in a physiologically relevant environment (**Figure** 7B, **S16**B). Next, we conducted electrophysiological recordings using PolyGraph microneedle electrodes, with classical borosilicate microneedles as control electrodes (Figure S16A). PolyGraph microneedles captured electrical activity, encouraged by potassium-rich artificial cerebrospinal fluid (high-K aCSF) perfusion, in murine brain slices, with waveforms characteristic of extracellular neuronal events (Figure 7C, 7E, S17).

Signal-to-noise ratio (SNR) analysis revealed enhanced performance compared to standard pipette electrodes (Figure 7D), and cross-correlation confirmed consistent detection of matching neuronal activity with minimal temporal lag, due to spatial separation (**Figure** 7F, **S18**). Recorded events exhibited diverse amplitudes and durations (**Figure** 7G, 7H, **S19, S20**), reflecting physiological events such as population spikes from groups of pyramidal neurons, field excitatory postsynaptic potentials (fEPSPs) and other network dynamics.^125–127^

To extract additional physiological insights, *k*-means clustering was applied to recorded event features to classify them according to their shape. This revealed three distinct waveform groups, visualised by their representative average traces with variability bands (**Figure** 7I, **S21**, **S22**). These clusters likely represent events from a mixture of different origins (e.g. population spiking, fEPSPs, fIPSPs, etc.), that enclose various phenomena (population synchronicity, excitation/inhibition balance, etc.), offering a valuable layer of physiological context for brain-computer interface (BCI) applications.

With a view to achieving on-chip event classification, a simple multi-layer perceptron and linear transformer classifier were trained on the clustered events. Both models achieved high classification accuracy (Figure 7J), enabling automated, real-time discrimination of neuronal events, allowing for interpretation and closed-loop modulation of neuronal signalling in future devices for both BCI applications and the treatment of diseases such as epilepsy and Parkinson’s disease.^9,10^

These results demonstrate PolyGraph10%’s capacity for high-resolution, bidirectional interfacing, showcasing current delivery, neuronal activity recording, and machine learning-based classification. This establishes PolyGraph10% as a next-generation neural interfacing platform, for closed-loop neuromodulation and real-time BCI applications.

Building on PolyGraph as a neural interfacing platform material, several avenues for future development remain. Chronic *in vivo* studies will be critical to evaluate long-term biocompatibility and integration with brain tissue, mechanical stability, and high-resolution recording performance in physiological environments.^128–130^ Leveraging alternative polymer matrices such as shape-memory, stimuli-responsive, elastomeric, and hydrogel materials will expand the range of accessible applications, and unlock complex device designs such as neural meshes and threads.^18,131,132^ Incorporating biohybrid strategies such as cell-laden hydrogel coatings could further mitigate immune responses and enhance tissue integration.^54^ Alternative surface treatments, including conductive polymer coatings (e.g., PEDOT:PSS)^133,134^ and nanoscale texturing,^135–137^ may extend electrochemical performance for both stimulation and recording applications. At the device level, scaling to high-density, multiplexed arrays with individually addressable channels across a wide area of coverage would enable next-generation closed-loop systems.^132^ Finally, the integration of machine learning-based classification demonstrated here offers a promising route towards real-time, on-chip analytics, showing that these advanced materials are compatible with state-of-the-art signal processing and analysis techniques for BCIs. Together, these advances stand to position PolyGraph as a transformative platform for minimally invasive, chronic neural interfacing across both the central and peripheral nervous systems, with far-reaching implications for clinical and neurotechnological applications.

## 3 Conclusions

The pursuit of soft, flexible, high-performance neural interfaces demands electroconductive materials that balance biocompatibility, stability, and low neuroinflammatory potential with electrical performance, processability, and flexibility. This work introduces PolyGraph, a novel PVP-stabilised graphene-polycaprolactone nanocomposite optimised for neural interfacing, combining flexibility, excellent electrochemical performance, and physiologically relevant biocompatibility. Surface engineering via NaOH roughening and AuPd coating yielded low impedance (∼1.6 Ω cm^2^ @ 1 kHz) and high charge injection capacity (11.7 mC/cm^2^ for a 100 ms pulse), key benchmarks for neural stimulation and recording. These composites were cast and 3D printed into freestanding microneedle arrays, connected using a bioresorbable hyaluronic acid-based backing to form a soft, multichannel electrode for minimally invasive, high-resolution neural interfacing. Bidirectional functionality was validated *ex vivo*, with PolyGraph10% microneedles delivering controlled stimulation and recording of extracellular neuronal activity from murine brain slices, alongside machine learning-based classification of neuronal events. Taken as a whole, these advances position PolyGraph10% as a versatile platform for soft bioelectronics, poised to drive progress in neuroprosthetics, regenerative medicine, and next-generation brain-computer interfaces.

## 4 Methods

### 4.1 Exfoliation of Graphene

#### 4.1.1 Gelatin graphene exfoliation - GelGr

Bovine gelatin (30 mg/mL, Sigma-Aldrich, Ireland) was dissolved in deionised (DI) water, and mixed with graphite (50 mg*/*mL, Graphexel Ltd, UK). The dispersion was exfoliated by probe ultrasonication (VCX-750, Sonics, USA, amplitude 80%, max power 200 W, cycle 6 s on/2 s off) for 72 h at room temperature (RT). Liquid cascade centrifugation (LCC)^138^ at 500 rpm and 1000 rpm removed unexfoliated graphite and obtain size selected graphene. The dispersion was then repeatedly washed at 2000 rpm, redispersing in DI water each time until gelatinous bubbles no longer formed upon shaking. The final concentration was then determined by vacuum filtration.

#### 4.1.2 Trypsin digestion of gelatin graphene - TrypGr

To enhance conductivity by thinning the gelatin coating on graphene, 20 mL of gelatin-graphene and 10 mL of trypsin solution (Sigma-Aldrich, Ireland) were stirred on a hot plate at 50 °C for 8 h. The resulting dispersion was centrifuged at 6000 rpm for 90 min, and the sediment was washed twice in DI water at 6000 rpm to remove excess trypsin and peptides.

#### 4.1.3 PVP (polyvinylpyrrolidone) graphene exfoliation - PVPGr

Graphite (40 mg/mL, Graphexel Ltd, UK) was dispersed in 80 mL isopropanol (99.9% HPLC IPA, Sigma-Aldrich, Ireland) with 2 mg*/*mL of PVP powder (Sigma-Aldrich, Ireland). The dispersion was then sonicated (VCX-750, Sonics, USA) for 12 h at 55% amplitude, on a 6 s on/2 s off cycle. Size selection was carried out by LCC between 1000 and 6000 rpm. To remove excess PVP or exchange solvents, the dispersion was centrifuged four times at 6000 rpm for 90 min. The above exfoliation parameters were varied to determine the optimal parameters for conductivity and yield of the PVPGr exfoliation:

**Methods Table 1.** Exfoliation parameters for PVP graphene optimisation testing: Description of exfoliation parameters tested during optimisation of polyvinylpyrrolidone (PVP) based exfoliation protocol.

| Method | PVP Concentration (mg/mL) | Graphite Concentration (mg/mL) | Solvent | Sonation Time (h) | Sonation Power (W) |
| --- | --- | --- | --- | --- | --- |
| PVPGr A | 2 | 50 | IPA | 72 | 250 |
| PVPGr B | 2 | 50 | IPA | 8 | 375 |
| PVPGr C | 2 | 50 | IPA | 12 | 375 |

#### 4.1.4 Surfactant exfoliation of graphene - SCGr

Graphite (40 mg/mL) was dispersed in 80 mL DI water with sodium cholate surfactant (6 mg/mL, SC, Sigma-Aldrich, Ireland). To pretreat the dispersion and remove impurities, the dispersion was probe ultrasonicated for 1 h, (55% amplitude, max power 750 W, cycle of 6 s on/2 s off). The dispersion was then centrifuged at 6000 rpm for 60 min, and the supernatant was discarded. The sediment was redispersed in 80 mL DI water, and 2 mg*/*mL SC was added. The dispersion was then sonicated for 8 h at 55% amplitude, on a 6 s on/2 s off cycle. Size selection was carried out by LCC between 1000 and 6000 rpm. To remove excess SC or exchange solvents, the dispersion was centrifuged four times at 6000 rpm for 90 min.

### 4.2 Physical Characterisation of Graphene

#### 4.2.1 Scanning Electron Microscopy (SEM) to assess material and device morphology

Samples were sputter coated with a 5 nm layer of an 80:20 gold-palladium mixture using a Cressington 108 auto sputter coater. Imaging at varying magnifications was performed on a Zeiss Ultra FE-SEM using the InLens detector at 3 kV accelerating voltage, 30 µm aperture size, and 5 mm working distance.

#### 4.2.2 Thermogravimetric analysis (TGA) to assess degree of polymer coating on graphene

TGA was performed using a Q50 thermogravimetric analyser (TA Instruments, USA). Samples were heated in air at 10 °C min^−1^, from 25 °C to 1000 °C.

#### 4.2.3 Ultra-violet/Visible Light (UV-Vis) Spectroscopy to investigate optical properties of graphene dispersion

UV-Vis spectroscopy was conducted between 220 and 1600 nm using a Perkin-Elmer Lambda 1050+ photospectrometer with an integrating sphere, to collect scattering, absorption, and extinction spectra.

#### 4.2.4 Raman Spectroscopy to investigate graphene defects and lateral size

IPA-graphene dispersions were dried onto silicon substrates at 100 °C. Raman spectra were acquired using a WITec RISE Raman system with a 633 nm laser, under ambient conditions with a spectral resolution of 2.5 cm^-1^ using a 600 g/mm grating.

#### 4.2.5 Atomic Force Microscopy (AFM) to assess graphene dimensions

AFM was performed on a Bruker Multimode 8 microscope. Diluted graphene inks (1:100 in IPA) were drop-cast onto Si/SiO_2_ substrates and imaged using OLTESPA R3 cantilevers in ScanAsyst mode. Statistical data were derived from the analysis of 173 flakes using Gwyddion SPM software.^139^ The lateral dimensions were calculated by computing the square root of the product of the flake length and width.

#### 4.2.6 Contact Angle measurements to assess hydrophilicity of nanomaterial composites

DI water contact angles were measured on 10 mm PVPGr/PCL discs using an OCA 25 contact angle goniometer (DataPhysics Instruments, USA), dispensing 5 µL DI water per drop.

#### 4.2.7 Mechanical Testing to measure stress-strain curves and buckling strength

Mechanical properties were derived from stress-strain curves obtained using a Zwick Z0.5 Proline tensile tester fitted with 5 N and 100 N load cells, at a strain rate of 0.01 %*/*s. Samples strips were clamped and tested in uniaxial tension.

#### 4.2.8 Optical Profilometry to assess surface roughness of nanomaterial composites

3D surface profiles were obtained using a Profilm3D Optical Profiler (Filmetrics) in white-light interferometry (WLI) mode with a 50× Nikon DI objective lens. Calibration was performed using a gold thin film on Si/SiO_2_ with a 50 nm step height confirmed by AFM. Profiles were levelled using a three-point levelling method, and step heights were calculated with the histogram step-height tool. The profile was smoothed using the “remove outliers” function in ProfilmOnline.

### 4.3 PCL-Graphene Composite (PolyGraph) Manufacture

Graphene pellets of known mass were sedimented from IPA by centrifugation at 6000 rpm for 90 minutes. The supernatant was discarded and replaced with polycaprolactone (PCL, M_w_ = 25 kDa) dissolved in an organic solvent (dichloromethane or chloroform, 100 mg/mL). The composite slurry was then redispersed using a tapered probe ultrasonicator for 2 hours at 30% amplitude. This slurry was then cast and dried into thin layers in Teflon molds.

**Methods Figure 1.**
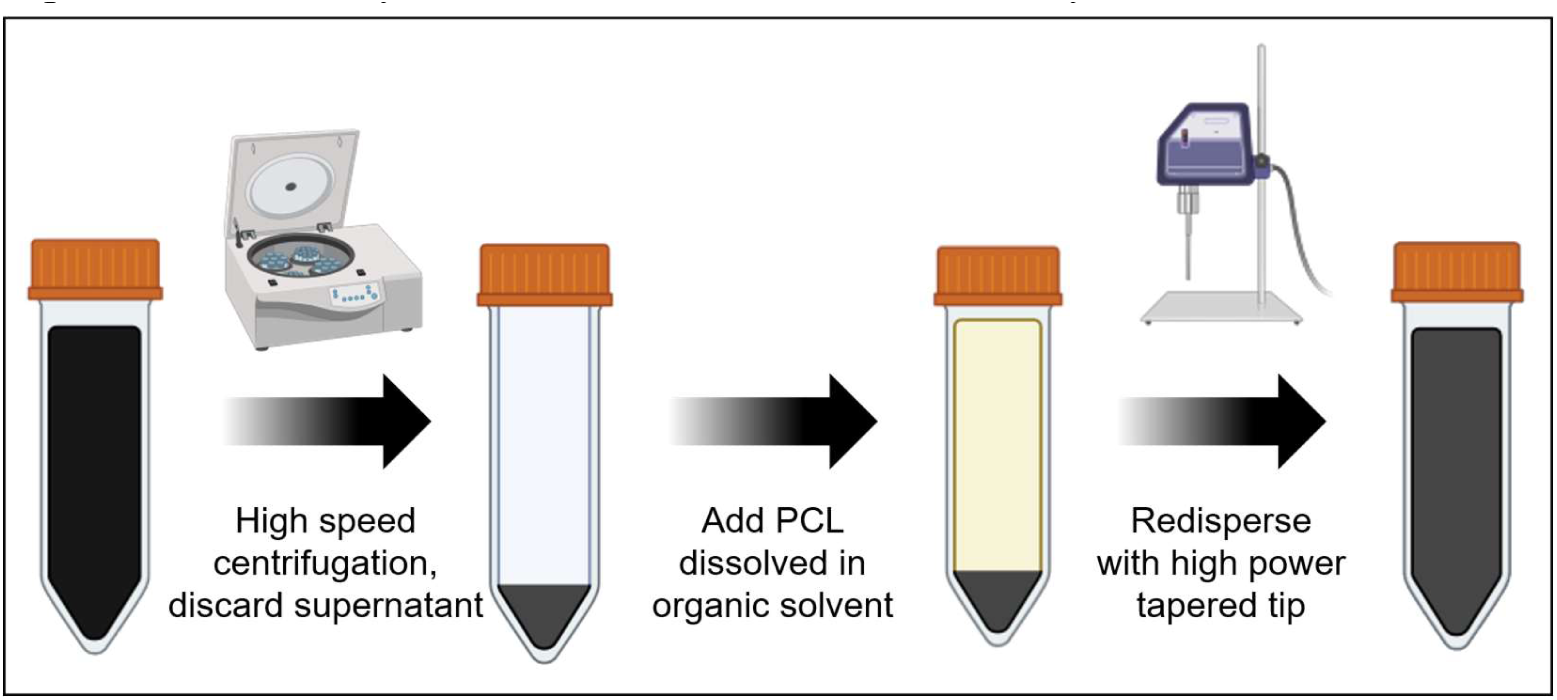
Fabrication of graphene-polymer composite slurry: Graphene dispersion of known concentration was sedimented by centrifugation at 6000 rpm. PCL, dissolved in an organic solvent, was added to achieve a 100 mg/mL slurry, which was thoroughly redispersed using a tapered sonic tip.

### 4.4 Cell Culture for Biocompatibility and Stimulation Testing

#### 4.4.1 NSC-34 mouse motor neurons

NSC-34 cells (Cedarlane Laboratories) were cultured in Dulbecco’s Modified Eagle Medium (DMEM, Sigma-Aldrich) supplemented with 10% fetal bovine serum (FBS, Biosera, Ireland), 1% Penicillin/Streptomycin (P/s, ScienCell, USA) and 4 mM L-glutamine (Sigma-Aldrich, USA).

#### 4.4.2 SH-SY5Y human neuroblastoma cells

SH-SY5Y cells (CRL-2266, ATCC) were cultured in a 1:1 mixture of Eagle’s Minimum Essential Medium and F12 Medium (Sigma-Aldrich, Ireland), supplemented with 10% FBS, 1% P/s, and 4 mM L-glutamine. For neuronal differentiation, cultures were switched to Neurobasal medium (Gibco, UK) with 2% B27 (Gibco, UK), 1% GlutaMAX (Sigma-Aldrich, Ireland) and 26.5 μM all-trans retinoic acid (atRA, Sigma-Aldrich, Ireland).

#### 4.4.3 iPSC-derived neuronal cells

iPSC neural precursor cells differentiated into neurons over 35 days as described previously.^140^ Cells were cultured in neural maintenance media (NMM): DMEM/F12 & Neurobasal medium (1:1), 1% P/s, 1% GlutaMAX, 1% MEM non-essential amino acids (NEAA), 0.2% N21 (Biotechne, UK), 1% B27 and 75 µM 2-mercaptoethanol (ThermoFisher, UK). For passaging, cells were detached using accutase (4-6 min, RT), centrifuged (1000 rpm, 3 min), and replated on Celltrex-coated (1:100, R&D Systems, UK) 6-well plates.

#### 4.4.4 iPSC-derived astrocytes

iPSC astrocyte progenitor cells^140^ were differentiated into astrocytes over 80 days using established protocols.^141,142^ Cells were cultured in Serio EL medium,^143^ consisting of Advanced DMEM-F12 (Gibco, UK), with 1% P/s, 1% GlutaMAX, 1% NEAA, 1% N21, 0.2% B27, 20 ng/mL human epidermal growth factor (EGF) and 20 ng/mL human leukaemia inhibitory factor (LIF) (Peprotech, UK).

#### 4.4.5 Primary neurons

E18 mouse cortices were removed post-mortem (Ethics Approval REC202005013) and placed in cold 0.1 M PBS. Cells were dissociated in a solution containing papain (40 U/mL, Worthington, USA), deoxyribonuclease I (100 U/mL, Worthington, USA), and 30 mM glucose in Hank’s Balanced Salt Solution (Sigma-Aldrich, Ireland), for 45 minutes at 37°C, 5% CO_2_. 1 mL of plating media (Neurobasal Plus, 2% B27 Plus, 0.5 mM GlutaMAX, and 10% Heat Inactivated Horse Serum (Sigma-Aldrich, Ireland)) was then added. The tissue was dissociated by gentle trituration with fire-polished Pasteur pipettes (Fisher Scientific). Neurons were counted using a haemocytometer and seeded at 5×10^5^ cells per droplet on each film, followed by overnight incubation at 37 °C and 5% CO_2_. The next day, wells were flooded with 3 mL of complete media (plating media without horse serum), and media was changed every two days (1 mL per well).

### 4.5 Biocompatibility Assessment in 2D Culture

#### 4.5.1 Cells grown with nanomaterial in suspension with culture media

Graphene dispersions in IPA were centrifuged at 6000 rpm for 90 minutes, and the supernatant was replaced by fresh deionised water. This step was then repeated twice to remove any residual IPA, followed by sterilisation via autoclaving. Treatment media (40 and 80 µg/mL) was prepared in the relevant cell culture media. Cells were seeded at 2×10^4^ cells/well in growth media and cultured for 24 h before replacing it with treatment media containing graphene. Media replacement was carried out every 3 days.

#### 4.5.2 Cells grown on films of nanomaterial composite

10 mm PCL-graphene discs mounted in Cell Crowns (Scaffdex, Sigma-Aldrich, Ireland) were sterilised using ethanol, UV exposure (30 min), and dip-washing in 10% P/s. Cells were seeded at 1×10^4^ cells*/*film in a small drop of growth media and allowed to attach for 1 h at 37 °C, before flooding the well with media. Media replacement was carried out every 3 days. Metabolic activity and DNA content were measured at designated timepoints following manufacturer protocols.

#### 4.5.3 Immunostaining to assess cellular morphology

At specific time points, samples were washed in PBS and fixed with either 10% formalin (Sigma-Aldrich, Ireland) or 4% paraformaldehyde (PFA, Fisher-Scientific, Ireland) for approximately 15 min at RT, followed by three PBS washes. To assess mature neuronal morphology, samples were incubated overnight at 4 °C with rabbit anti-β Tubulin III primary antibody (1:1000, Sigma-Aldrich) in PBS, with 0.1% Triton-X100 and 1% bovine serum albumin. The following day, after three PBS washes, samples were incubated with Alexa Fluor® 555 Goat Anti-Rabbit IgG (1:1000, Invitrogen, UK) for 2 h at RT. To visualise cytoskeletal structure of both neuronal and glial cells, samples were incubated overnight at 4 °C with fluorescein isothiocyanate (FITC)-labelled phalloidin (1:1000, Sigma-Aldrich, Ireland). After three PBS washes, nuclei were counterstained with DAPI (1:1000) in PBS with 0.1% Triton-X100, and samples mounted on slides using Fluoroshield™ (Sigma-Aldrich). Imaging was performed using a Zeiss AxioObserver fluorescent microscope (Carl Zeiss Ltd., USA), and images analysed in FIJI (ImageJ 1.52p).

### 4.6 NaOH & AuPd Treatment to Enhance Electrochemical Behaviour of PolyGraph Composites

To enhance electrochemical performance, NaOH roughening was used to increase the specific surface area, while AuPd coating was used to reduce the surface impedance. Samples were submerged in 3 M NaOH aqueous solution for 72 h at RT, followed by rinsing with DI water. Samples were subsequently coated with a thin layer (∼10 nm) 80:20 gold-palladium mixture, using a Cressington 108 auto sputter coater.

### 4.7 Electrical Characterisation of Material and Device Performance

#### 4.7.1 Conductivity

DC conductivity was measured using a Keithley Model 2450 Sourcemeter, connected to silver paint electrodes on each end of thin strips of PCL-graphene.

#### 4.7.2 Cyclic Voltammetry (CV), Electrochemical Impedance Spectroscopy (EIS) & Chronoamperometry (CA) to measure Charge Storage Capacity (CSC), impedance, and electrochemical potential window

Two-electrode coin cells were fabricated with 3 mm diameter discs of PolyGraph, using phosphate-buffered saline (PBS) as the electrolyte, and polyethylene filters as separators (**Figure S10**A). CV and EIS measurements were performed on a Biologic VMP-300 potentiostat. CV was carried out in a ±200 mV potential window at scan rates from 1 mV/s - 30000 mV/s. EIS was recorded across 1 Hz – 7 MHz at 10 mV amplitude. CA was measured in steps of 100 mV from -1 V to 1 V, with the magnitude of the current transient at 1 s after stimulation extracted. Data analysis was carried out using Biologic EC-Lab software, to extract the surface impedance (*Z*), cut-off frequency (*f_cut-off_*), and charge storage capacity (CSC).

#### 4.7.3 Voltage Transients Analysis to measure Charge Injection Capacity (CIC)

Voltage transients were recorded using a Swagelok three-electrode system with glassy carbon electrodes, a graphite reference electrode, and a carbon black/Teflon counter electrode, using PBS as the electrolyte (**Figure S12**A). Chronopotentiometry delivered defined current densities, followed by measurement of the resulting voltage transients. The maximum polarisation at each current density was used to calculate the maximum charge injection capacity (CIC_max_) of the tested material.

### 4.8 Microneedle Array Manufacture for Flexible BCI Prototype

Microneedle bases were designed in AutoCAD Inventor, with the desired pitch, angle, and needle length. Designs were 3D printed in clear resin using an SLA printer (Form 3/4B, FormLabs, USA, then washed in IPA for 30 minutes, before 60 minutes of UV curing at 60 °C. Polydimethylsiloxane (PDMS, Sylgard 184, Ellsworth Adhesives, Ireland) was mixed, degassed, and poured over the microneedle bases in Petri dishes to form negative molds, which were then cured overnight at 60 °C. Negative molds were then carefully peeled from the microneedle base. This process was adapted from O’Cearbhaill et al.^144,145^

Microneedle arrays were fabricated in a multi-stage process (**Figure 6**A). PolyGraph films were placed in a microneedle mold at 200 °C until softened, followed by mechanical compression with a spatula, to penetrate all needle holes. For free-standing, non-isolated devices, the microneedle array was peeled from the mold at this step. To produce flexible, isolated devices, excess PolyGraph was removed while still molten. Pillars of PolyGraph, dissolved in dichloromethane at 1.5 g/mL, were printed on the back of the microneedles using custom G-code on an FDM printer (Allevi 3, Allevi, USA) and via manual extrusion using a syringe. A 5 mg/mL hyaluronic acid solution was then cast and dried between the pillars, to form a soft, conformable base. Final devices were gently released from the molds.

### 4.9 Assessment of Bidirectional Neural Interfacing Capabilities

#### 4.9.1 Ethical declaration for ex vivo experiments

All *ex vivo* mouse and rat tissues (mouse back tissue, rat spinal cord, and mouse brain slices), were obtained as post-mortem byproducts from ongoing experiments in other RCSI groups which have been approved by the RCSI Research Ethics Committee (REC 1587) and under license by the Health Products Regulatory Authority (AE19127/P057), in line with institutional animal welfare policy and the 3Rs (replacement, reduction, refinement).

#### 4.9.2 Recording of neuronal signals in brain slice model and delivery of stimulation pulses

Individual microneedle/pillar units were detached from the molds described in **Section 4.8**. These units were then NaOH and AuPd treated as described in **Section 4.6**, and connected to wires with silver paint. To reduce electrode size and increase recording resolution, the microelectrodes were sheathed in PDMS to just below the tip. Electrodes were wired to BNC connectors, and signals were pre-amplified (gain = 100, SR560, Stanford Research Systems, USA), digitized (Axon DigiData 1550B, Molecular Devices) and recorded via Clampex software (Molecular Devices) at 10 kHz, with low-pass filtering at 2 kHz. Analog signals were visualized on an oscilloscope (TBS1000C, Tektronix, USA). An AuPd-coated wire served as the reference electrode, and a glass electrophysiological pipette electrode served as the control.

Acute 350 µm thick coronal brain slices were obtained using a vibratome (7000smz-2, Campden Instruments, Loughborough, UK) from an 11-week-old C57/Bl6 male mouse. Briefly, the animal was deeply anesthetised (5% isoflurane) and quickly decapitated. The whole brain was subsequently removed and placed in ice-cold oxygenated (95% O_2_, 5% CO_2_) high-sucrose artificial cerebrospinal fluid solution (aCSF, 11 mM glucose, 1 mM MgSO_4_, 1 mM NaH_2_PO_4_, 2.5 mM KCl, 26.2 mM NaHCO_3_, 2.5 mM CaCl_2_, 119 mM NaCl and 75 mM sucrose), slices were then left to recover for at least 1 hour in a chamber containing Ringer solution (126 mM NaCl, 3 mM KCl, 1.25 mM NaH_2_PO_4,_ 2 mM MgSO_4_, 2 mM CaCl_2_, 10 mM glucose and 26 mM NaHCO_3_) at RT until recording.

After recovery (>1 hour), slices were transferred to a submerged recording chamber in a patch-clamp rig constantly perfused with oxygenated Ringer solution at physiological temperature (36 °C), at a flow rate of ∼7.5 mL/min. Under microscope visualization (Scientifica SliceScope), the microneedle and pipette electrodes were placed 10-100 μm apart on the stratum radiatum of the CA1 hippocampal region, and spontaneous neuronal activity was encouraged using a modified aCSF solution (0.4 mM MgSO_4_, 8 mM KCl, 300 µM picrotoxin (PTX)).

For stimulation, microneedle electrodes were connected to an IonOptix C-Pace stimulation rig delivering biphasic current pulses, with an AuPd-coated wire as counter electrode. The waveform of stimulation pulses was monitored on the oscilloscope via two additional AuPd-coated wires.

**Methods Figure 2.**
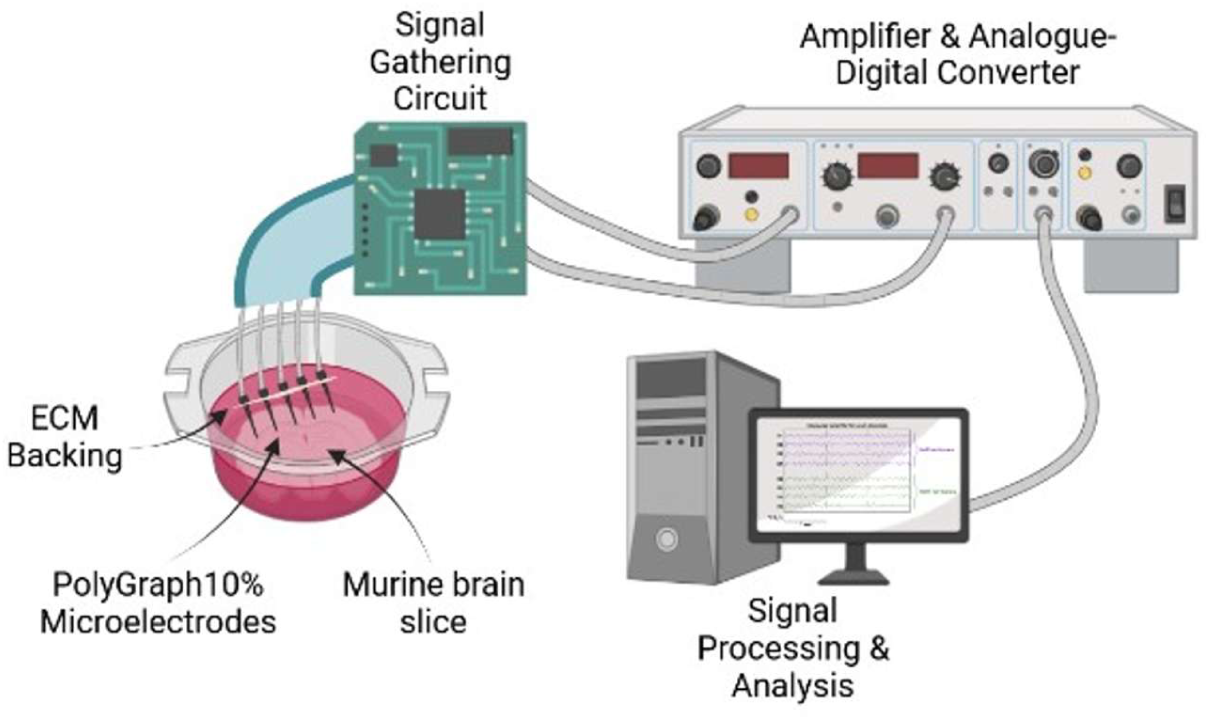
Bidirectional neural interfacing setup: Schematic of stimulation or recording from brain slice model using a flexible PolyGraph neural interface.

#### 4.9.3 Event analysis

Acquired recordings were notch-filtered at 50 Hz and its first five harmonics to remove electrical interference, and low-pass filtered at 150 Hz to remove high-frequency noise and isolate slower LFPs. A template event, manually identified from 100 traces, and used to identify 437 distinct events in the recording from the PolyGraph electrode. Traces were processed using a custom Python analysis pipeline, which aligns and scales the waveforms prior to extracting features such as the peak value, peak width, and area under curve.

Dimensionality reduction was performed using principal component analysis (PCA) and t-distributed Stochastic Neighbour Embedding (t-SNE), followed by *k*-means clustering. To test classification of unseen data, a multilayer perceptron (MLP) and linear transformer model were trained on a subset of the clustered data, and their accuracy evaluated using confusion matrices.

### 4.10 Statistical analysis

A minimum of three experimental replicates (N) with at least three technical replicates (n) were used in all assessments. Prior determination of statistical significance, the ROUT method (Q=10%) was used to determine the variance in the population. All data sets were subjected to Kolmogorov-Smirnov test to assess for normality and thereafter the data tested using appropriate parametric (t-test, one- and two-way ANOVA) and non-parametric (Mann-Whitney U, one and two-way Kruskall-Wallis) tests. Post-hoc tests were performed using Tukey’s multiple comparisons and Dunnett’s correction where needed. All results were plotted with the standard error of mean and were considered significant when p < 0.05 and statistical significance was denoted by *: *p < 0.05, **p < 0.01, ***p < 0.001, ****p < 0.0001. Significances are relative to control group of timepoint unless stated otherwise. All data were plotted and analysed as means with standard error of the mean while using the software GraphPad Prism (v8.0) and OriginLab 2024.

## Acknowledgements

The authors would like to acknowledge Prof. Oran D. Kennedy and Prof. Eoin O’Cearbhaill for insight into microneedle manufacture. Additionally, we would like to acknowledge the support given by the Advanced Microscopy Laboratory (AML, Trinity College Dublin) in the acquisition of SEM data. Finally, we would like to acknowledge the Department of Physiology, the FutureNeuro Centre, and Prof. David Henshall in RCSI for provision of electrophysiological recording infrastructure and donation of *ex vivo* animal tissue.

## 6 Data Availability Statement

The data that support the findings of this study are available from the corresponding author upon reasonable request.

## 8 Supporting Information

**Figure S1.**
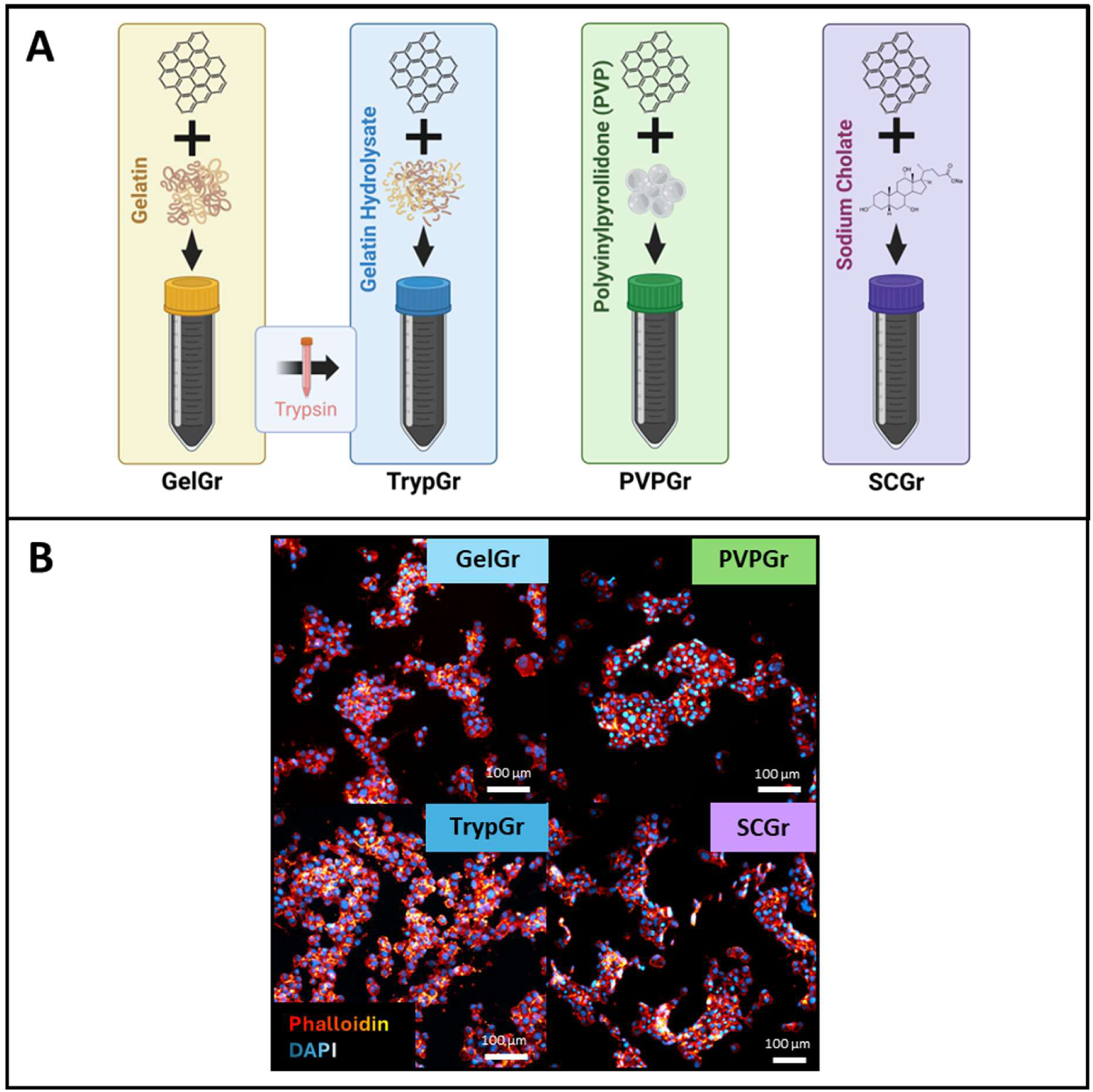
Biological characterisation of graphene candidates: **A)** Schematic of graphene formulations, highlighting stabilisation molecules used: gelatin (GelGr), gelatin hydrolysate post-trypsin treatment (TrypGr), polyvinylpyrrolidone (PVPGr), and sodium cholate (SCGr). **B)** Representative immunofluorescence images of cells cultured on collagen-graphene films of each formulation. Phalloidin: Yellow/red, DAPI: Blue/white. Scale bars 100 μm. Significances: *p < 0.05, **p < 0.01, ***p < 0.001, ****p < 0.0001.

**Figure S2.**
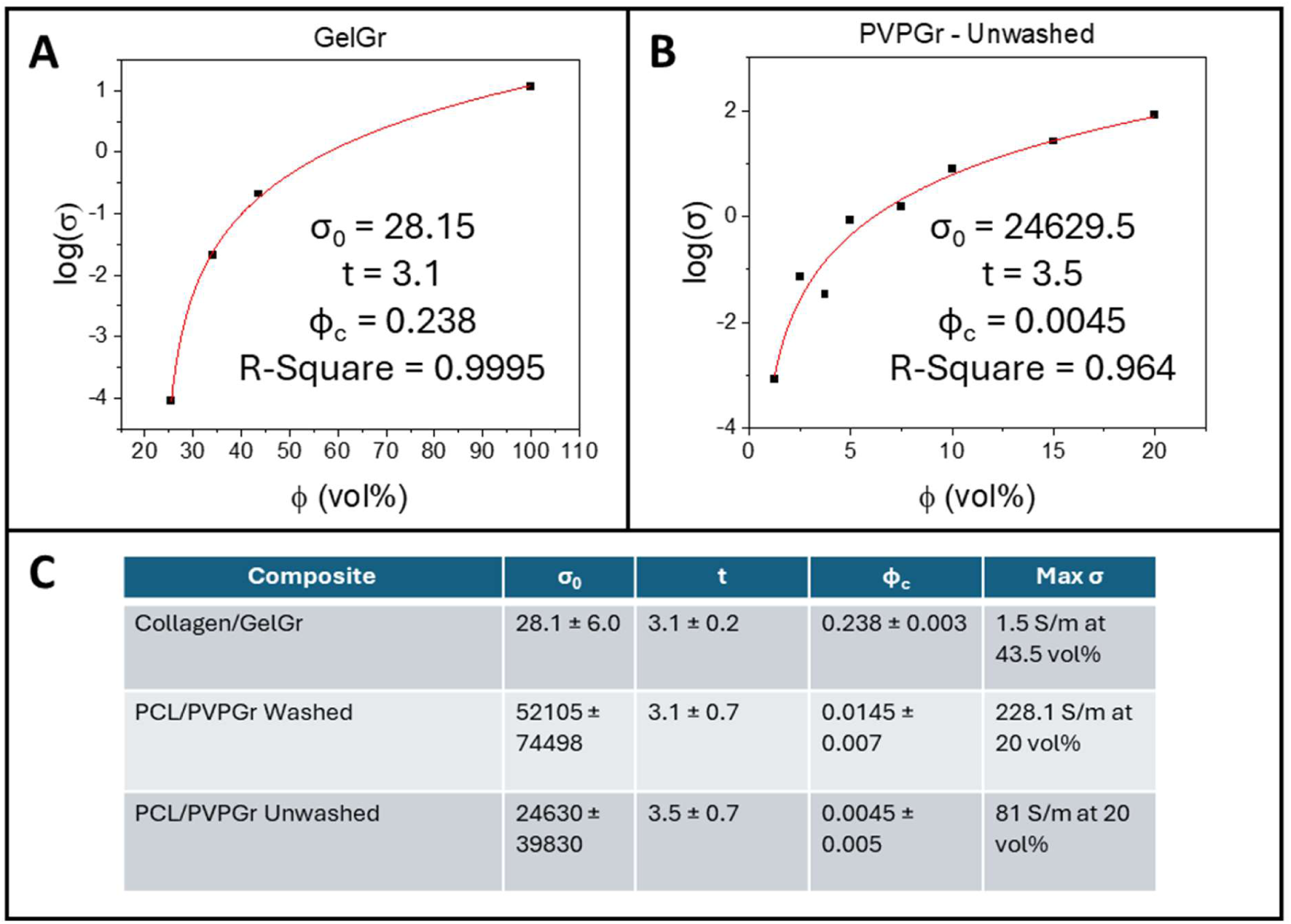
Percolation curves: A-B) Percolation curves for GelGr (A) and unwashed PVPGr (B). **C)** Summary of percolation fitting parameters (σ₀, t, ϕ_c_, and maximum σ), highlighting the superior conductivity of PolyGraph versus collagen/GelGr, and the critical role of washing in enhancing electrical properties.

**Figure S3.**
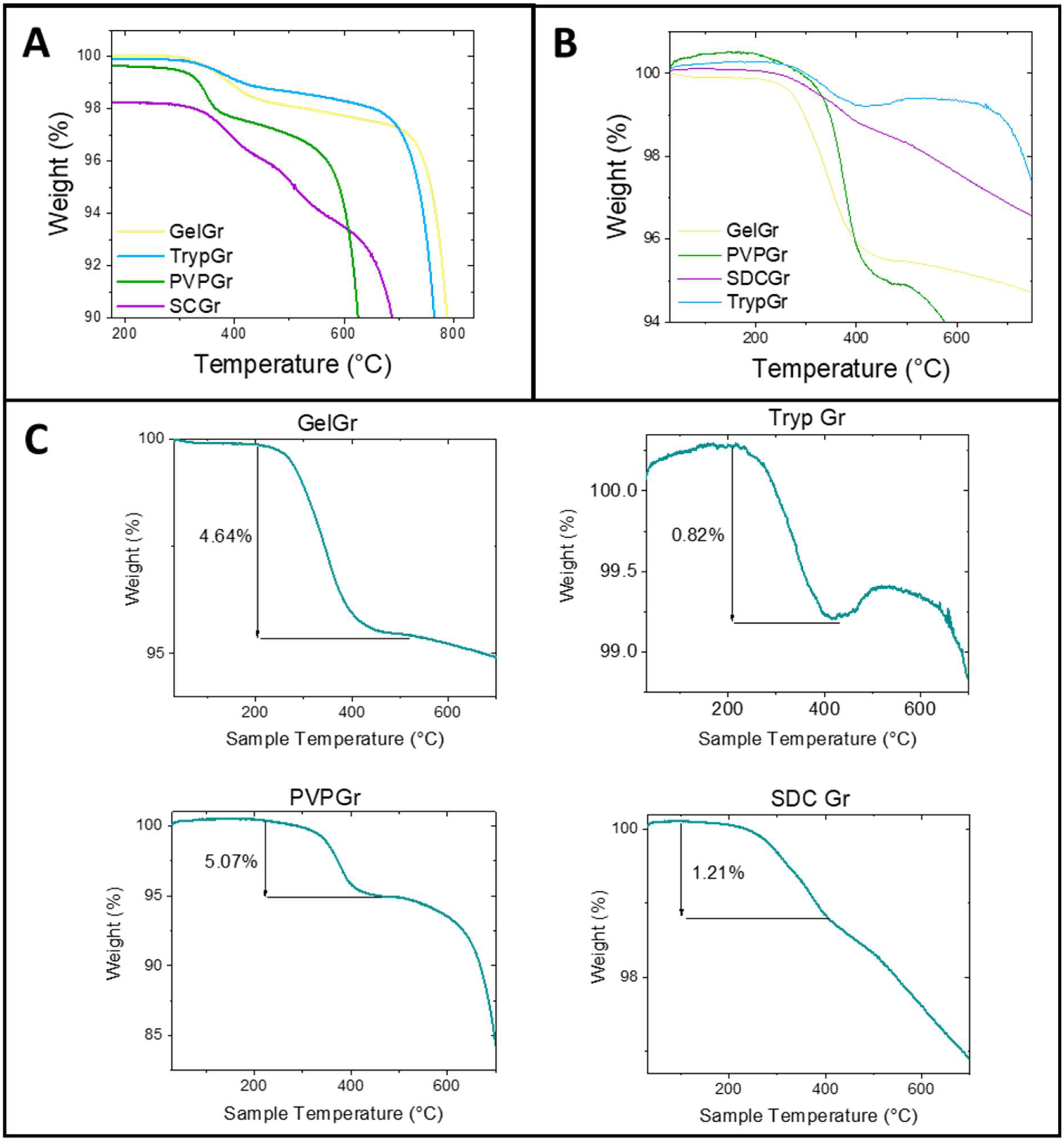
Thermogravimetric analysis (TGA) of graphene candidates: **A)** TGA curves in air, providing estimates of the surface coating thickness for each formulation. **B)** TGA performed under nitrogen atmosphere shows similar trends for GelGr, TrypGr, PVPGr, and SCGr. **C)** Individual TGA traces for each graphene type, highlighting percentage weight loss associated with polymer removal.

**Figure S4.**
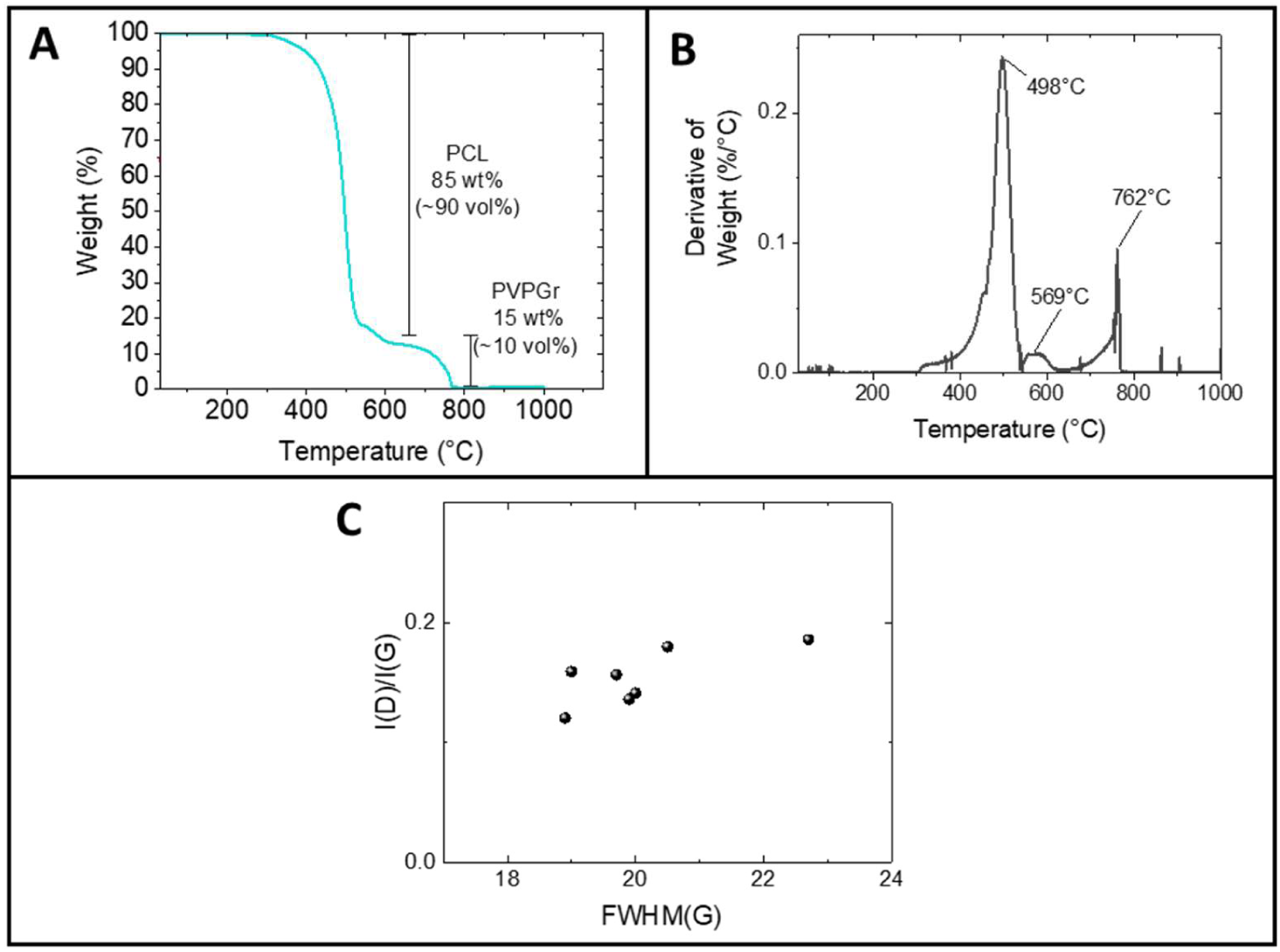
Further physical characterisation of PVPGr and PolyGraph: **A)** TGA of PolyGraph, verifying accurate formulation composition. **B)** Derivative of TGA spectrum for PolyGraph10%, showing decomposition peaks corresponding to PCL (∼500 °C) and PVPGr (∼750 °C) respectively. **C)** Raman analysis of PVPGr, showing no correlation between the I_D_/I_G_ ratio and the full-width-half-maximum (FWHM) of the G peak, indicating a defect-free basal plane.

**Figure S5.**
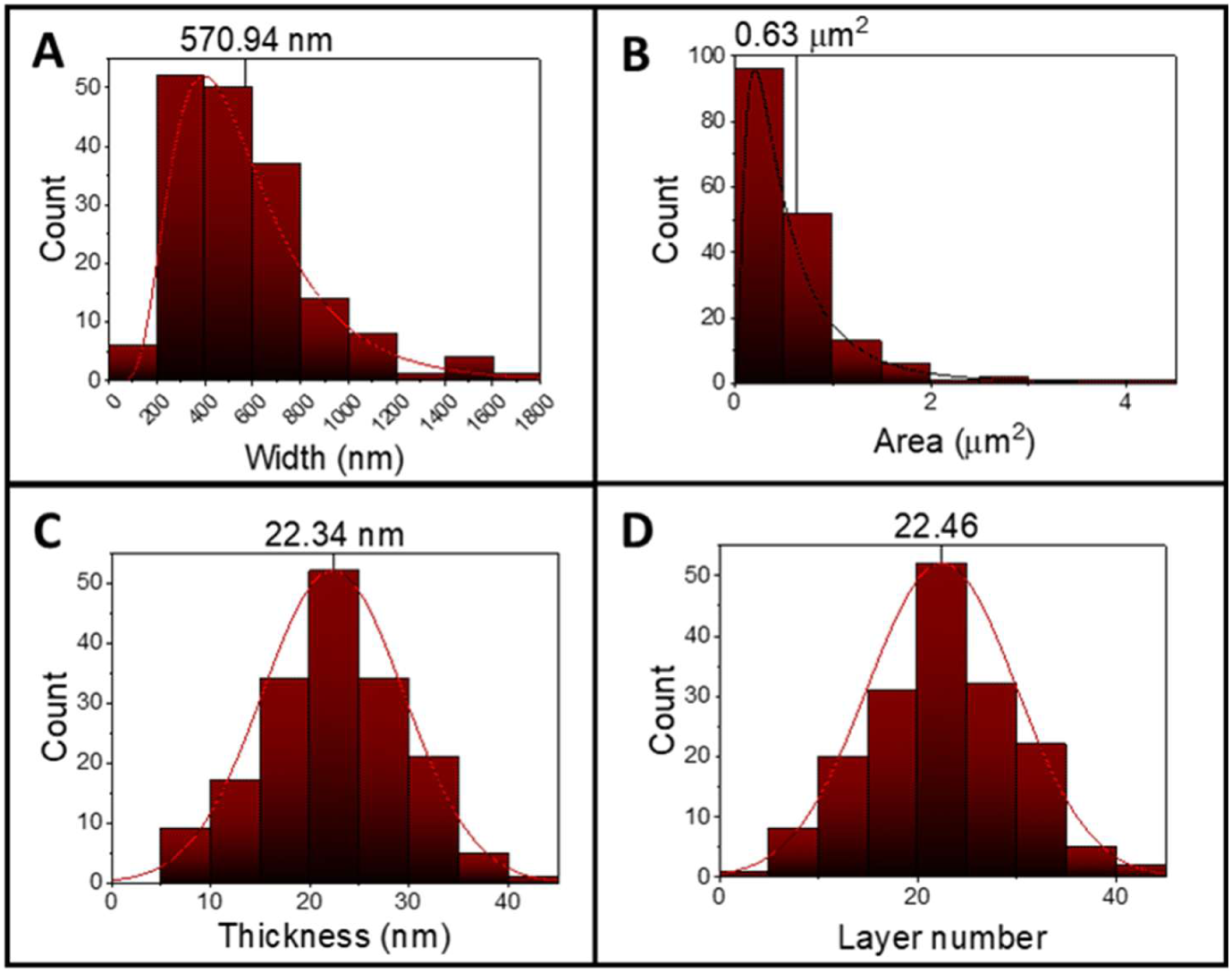
AFM analysis of PVPGr nanosheets: Histograms of **A)** nanosheet width **B)** nanosheet area **C)** nanosheet thickness and **D)** estimated layer number, based on analysis of 173 flakes. Mean values indicated above each plot.

**Figure S6.**
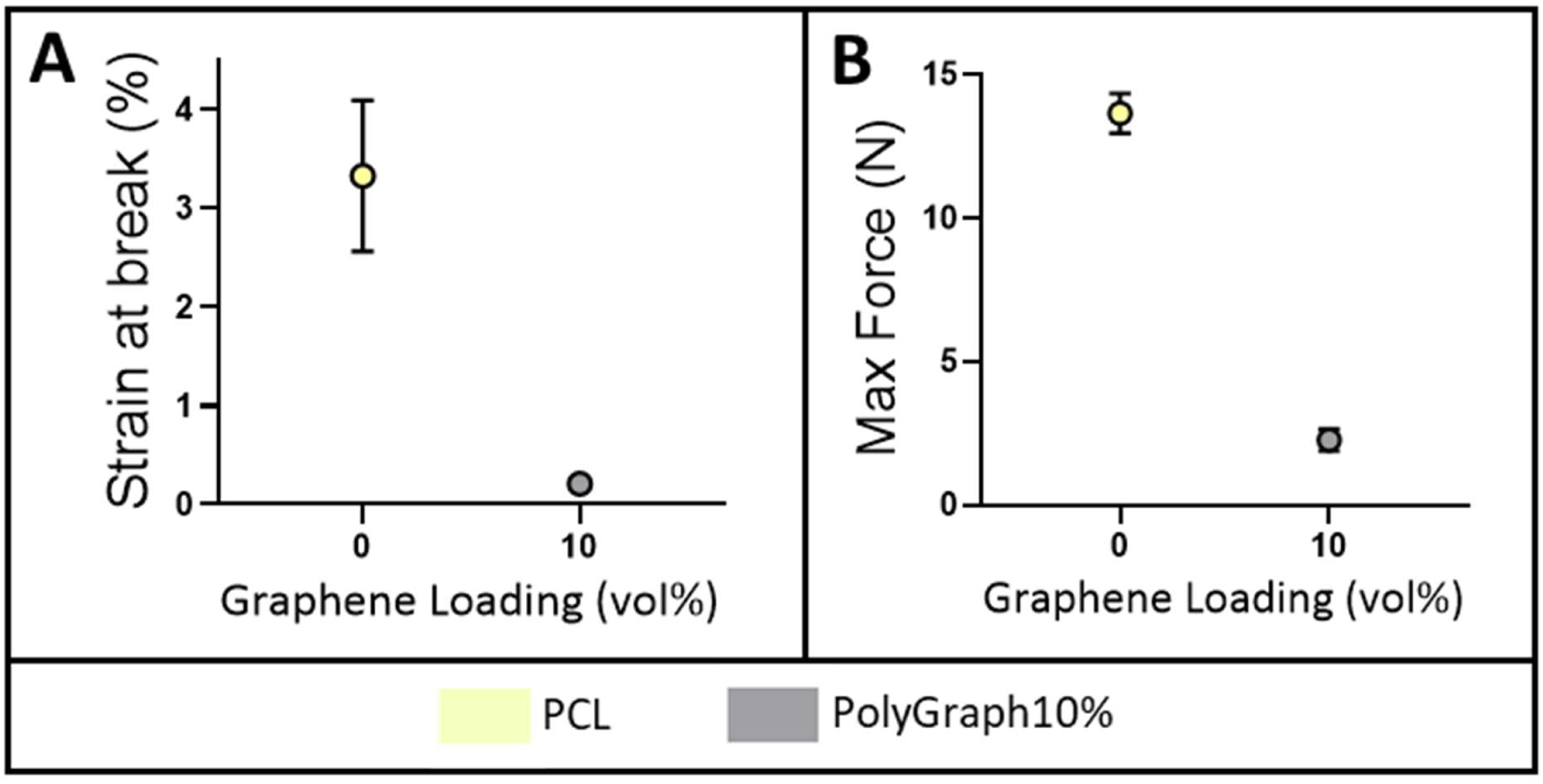
Mechanical testing of PolyGraph: A-B) Strain at break (A) and maximum force (B) measurements for PCL and PolyGraph10%, showing reduced ductility following graphene incorporation.

**Figure S7.**
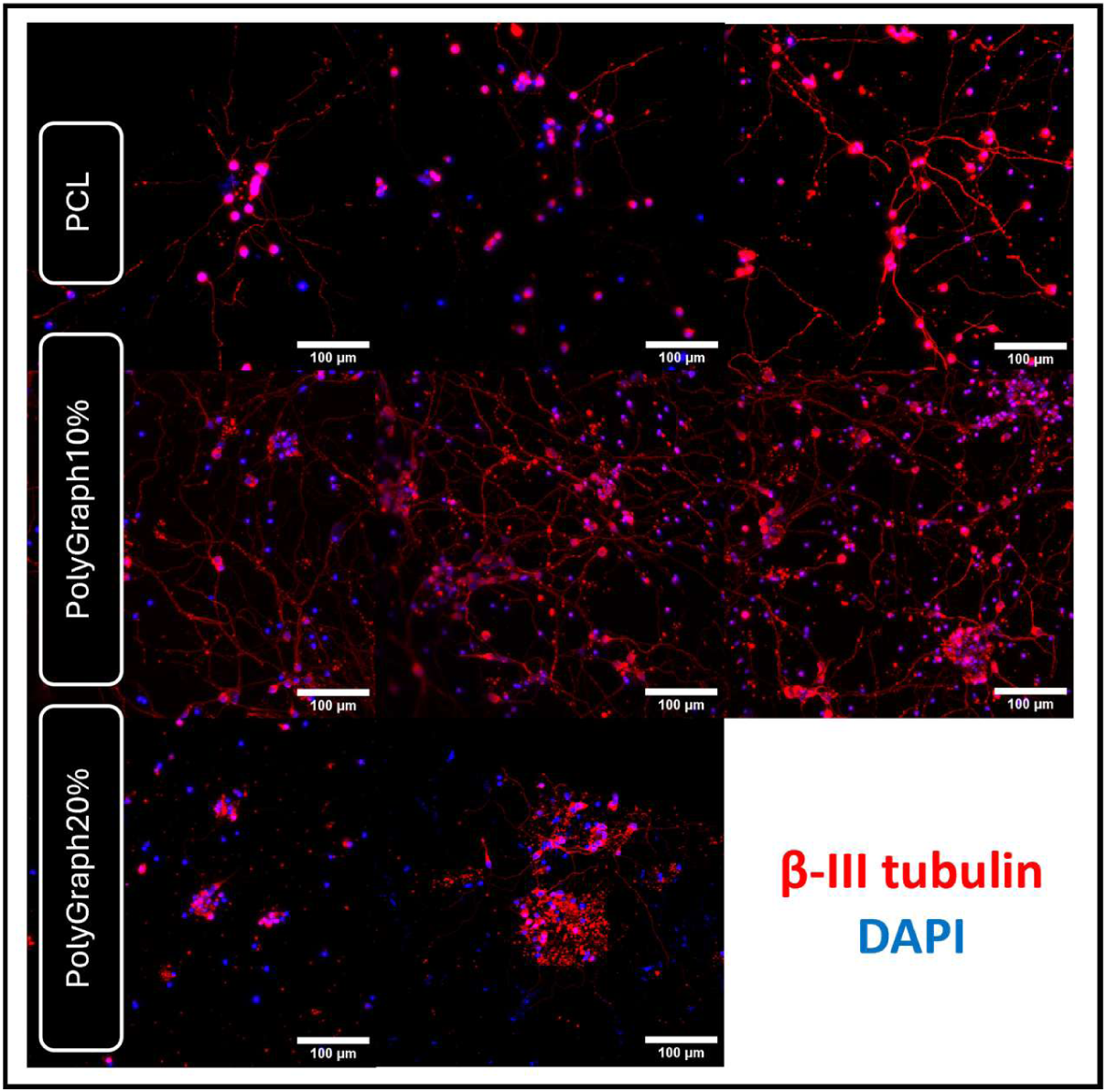
iPSC-derived neurons cultured on PolyGraph: iPSC-derived neurons were cultured on PolyGraph for 14 days and imaged using confocal microscopy. Extensive neurite outgrowth and network formation are visible on PolyGraph10%, indicating its support of neuronal growth. Scale bars 100 µm.

**Figure S8.**
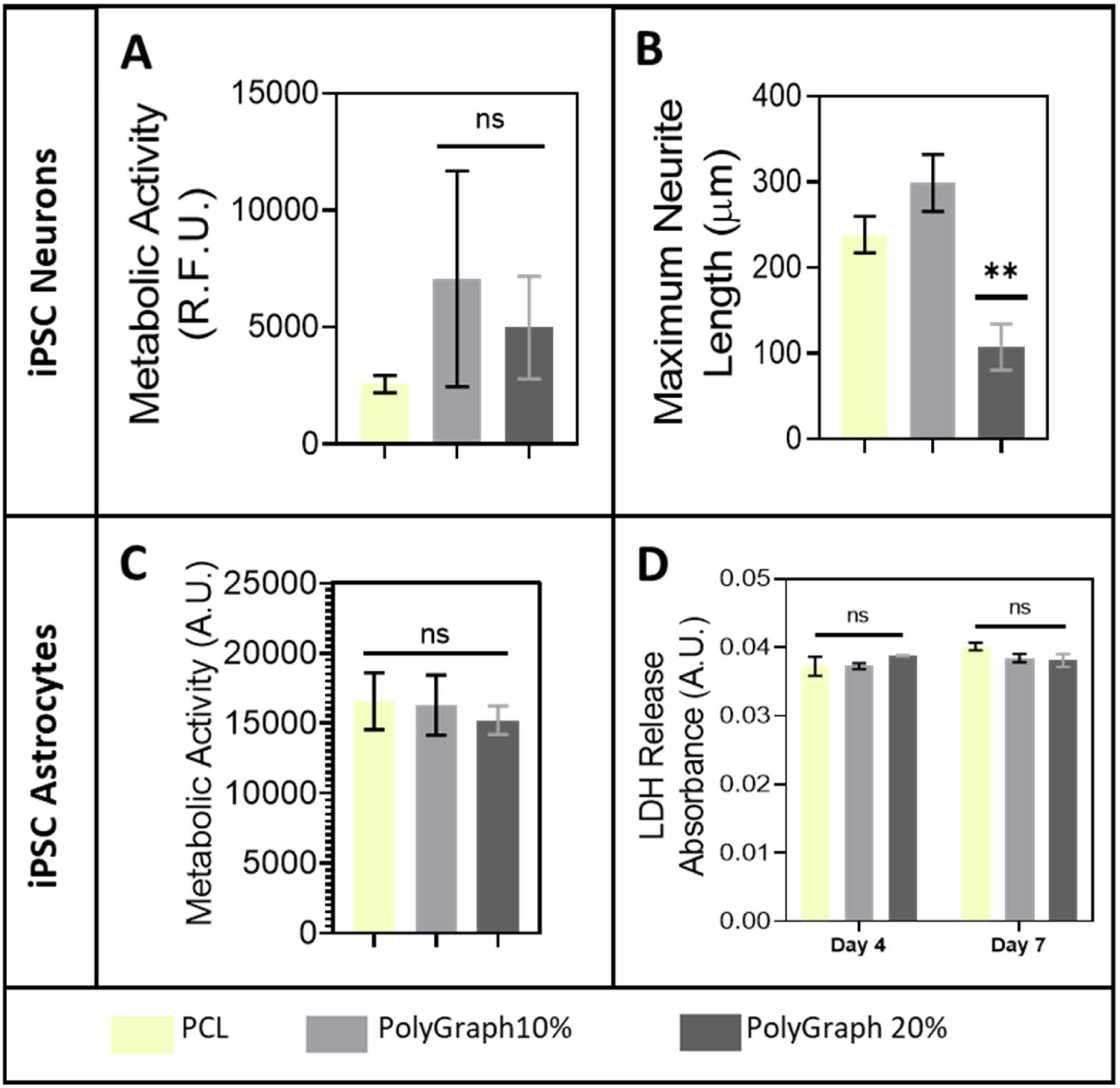
Biological and immunological assessment of PolyGraph: **A)** Metabolic activity of iPSC-derived neurons cultured on PolyGraph. **B)** Maximum neurite length of iPSC-derived neurons, showing a significant increase on PolyGraph10%. Scale bars 100 μm. **C)** LDH release from iPSC-derived astrocytes, indicating no significant cytotoxicity on PolyGraph. **D)** Metabolic activity of iPSC-derived astrocytes at day 14, showing no significant differences. Significances: **p < 0.01.

**Figure S9.**
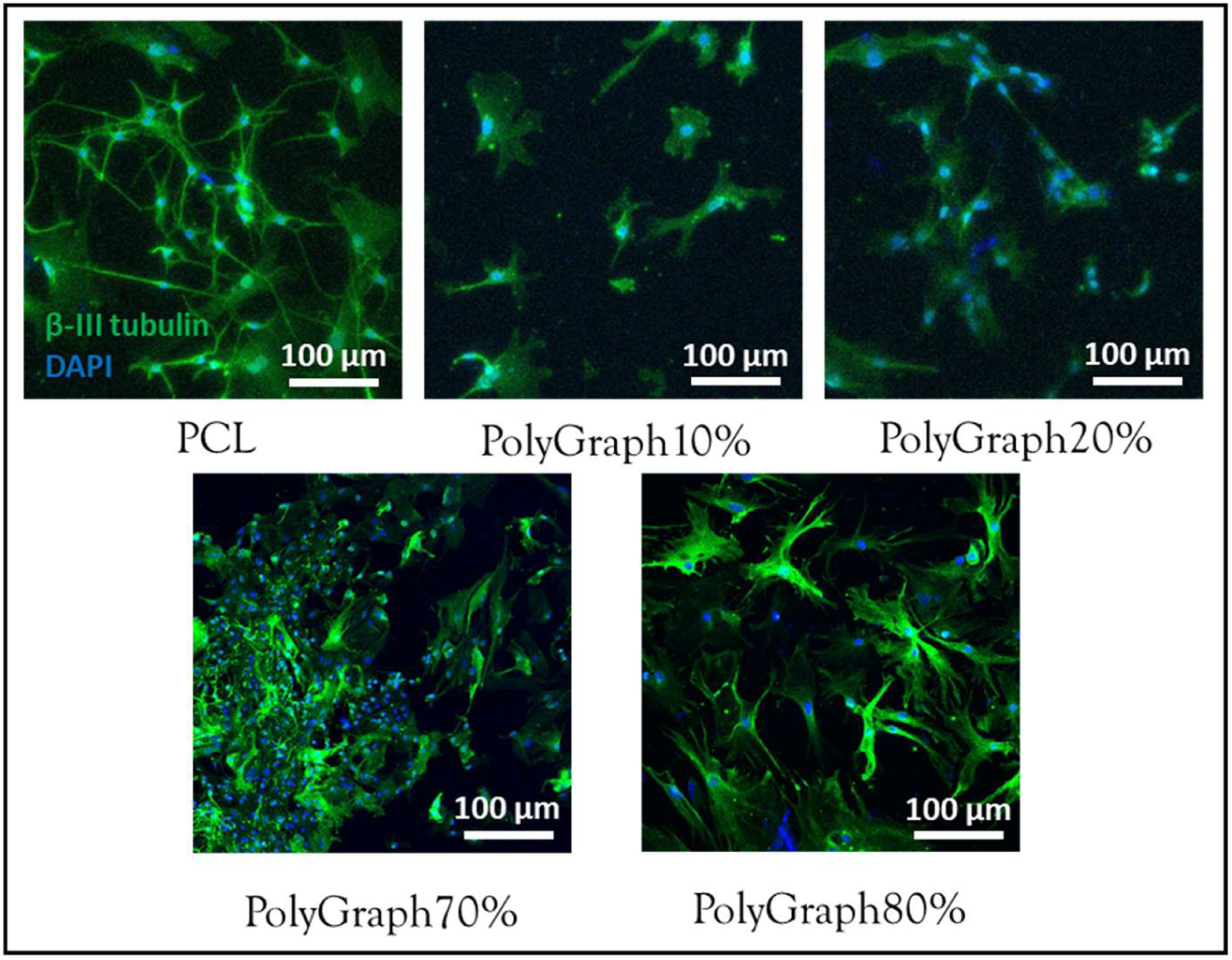
Mouse primary cortical neurons cultured on PolyGraph: Mouse primary neurons were cultured on PolyGraph composites for 14 days and imaged via confocal microscopy. Robust survival on PolyGraph composites indicates biocompatibility with sensitive, physiologically relevant cells. Scale bars 100 μm.

**Figure S10.**
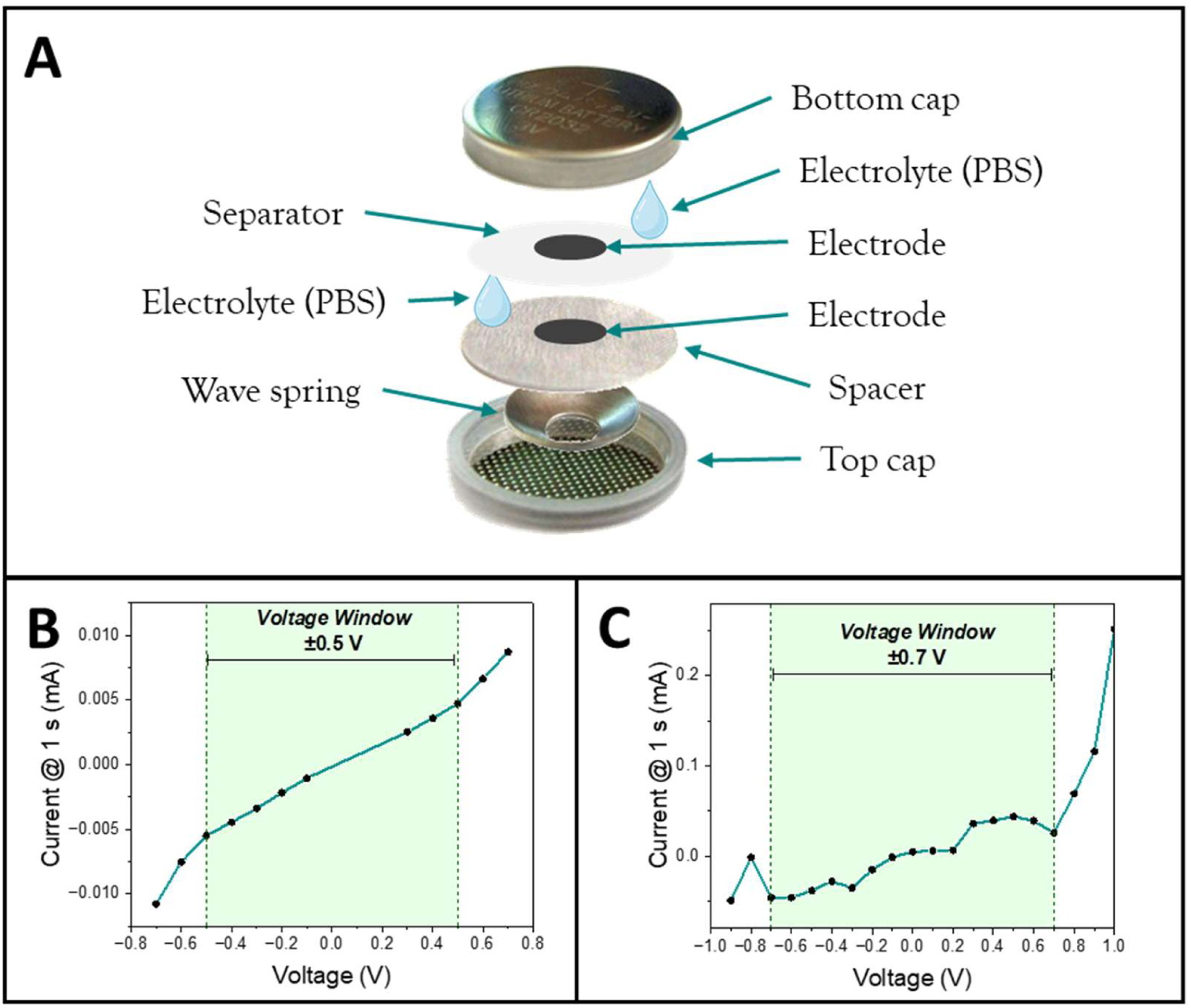
Electrical characterisation of PVPGr and PolyGraph composites: **A)** Schematic of two-electrode coin cell setup used for electrochemical characterisation. **B)** Chronoamperometry data showing the potential window of untreated PolyGraph at ±0.5 V. **C)** Chronoamperometry of AuPd + NaOH-treated PolyGraph, indicating an expanded potential window of approximately ±0.7 V.

**Figure S11.**
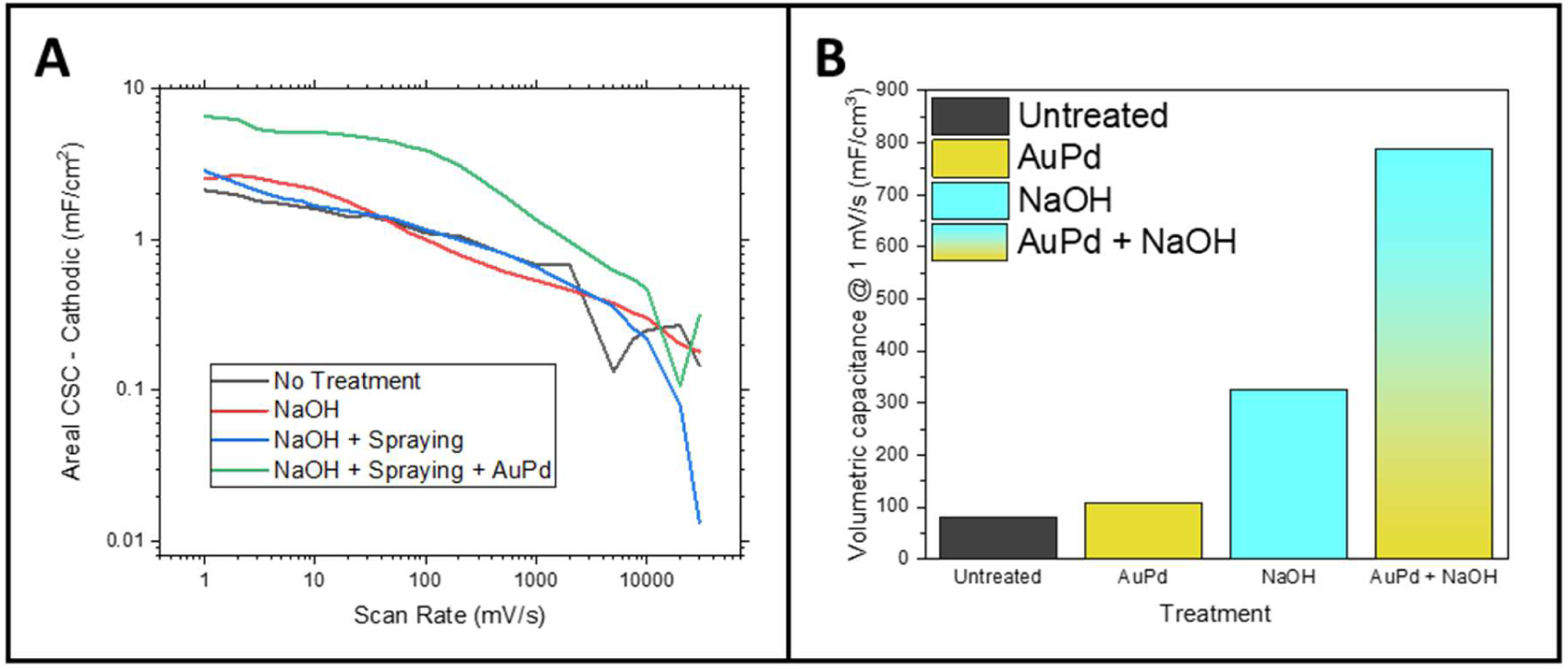
Charge storage capacity enhancement of PolyGraph by surface treatment: **A)** Areal charge storage capacity (CSC) of PolyGraph composites as a function of scan rate, showing the effects of NaOH roughening, graphene spray coating, and AuPd sputter coating individually and in combination. **B)** Volumetric capacitance at 1 mV/s for untreated and treated PolyGraph composites, highlighting substantial improvements with combined NaOH and AuPd treatments.

**Figure S12.**
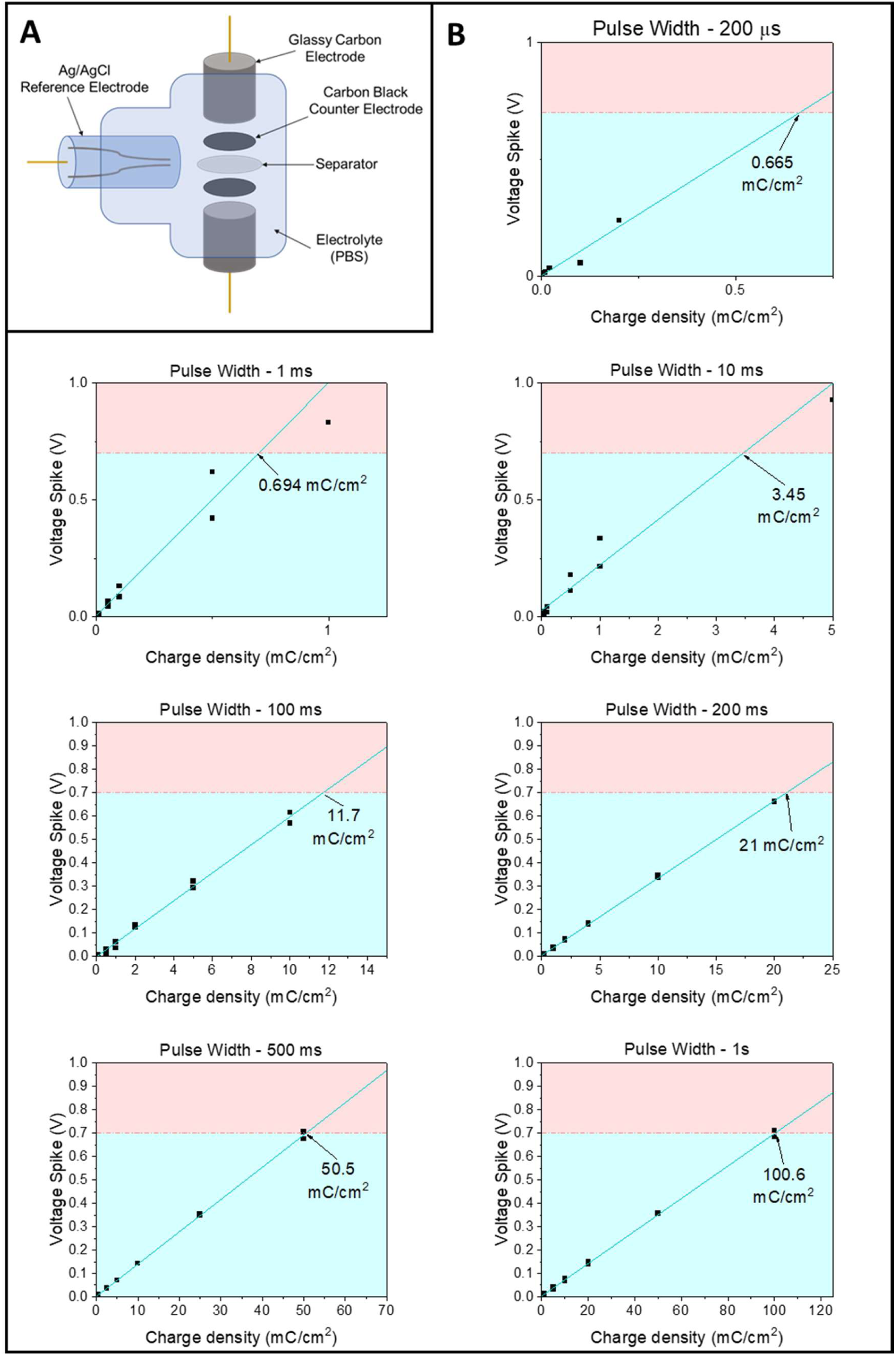
Voltage transients curves: **A)** Schematic of three-electrode setup used to capture voltage transients following square wave excitation **B)** Voltage transient responses at pulse widths from 200 µs to 1 s, used to calculate charge injection capacity (CIC). The electrochemical potential window (∼±0.7 V) is shown in blue.

**Figure S13.**
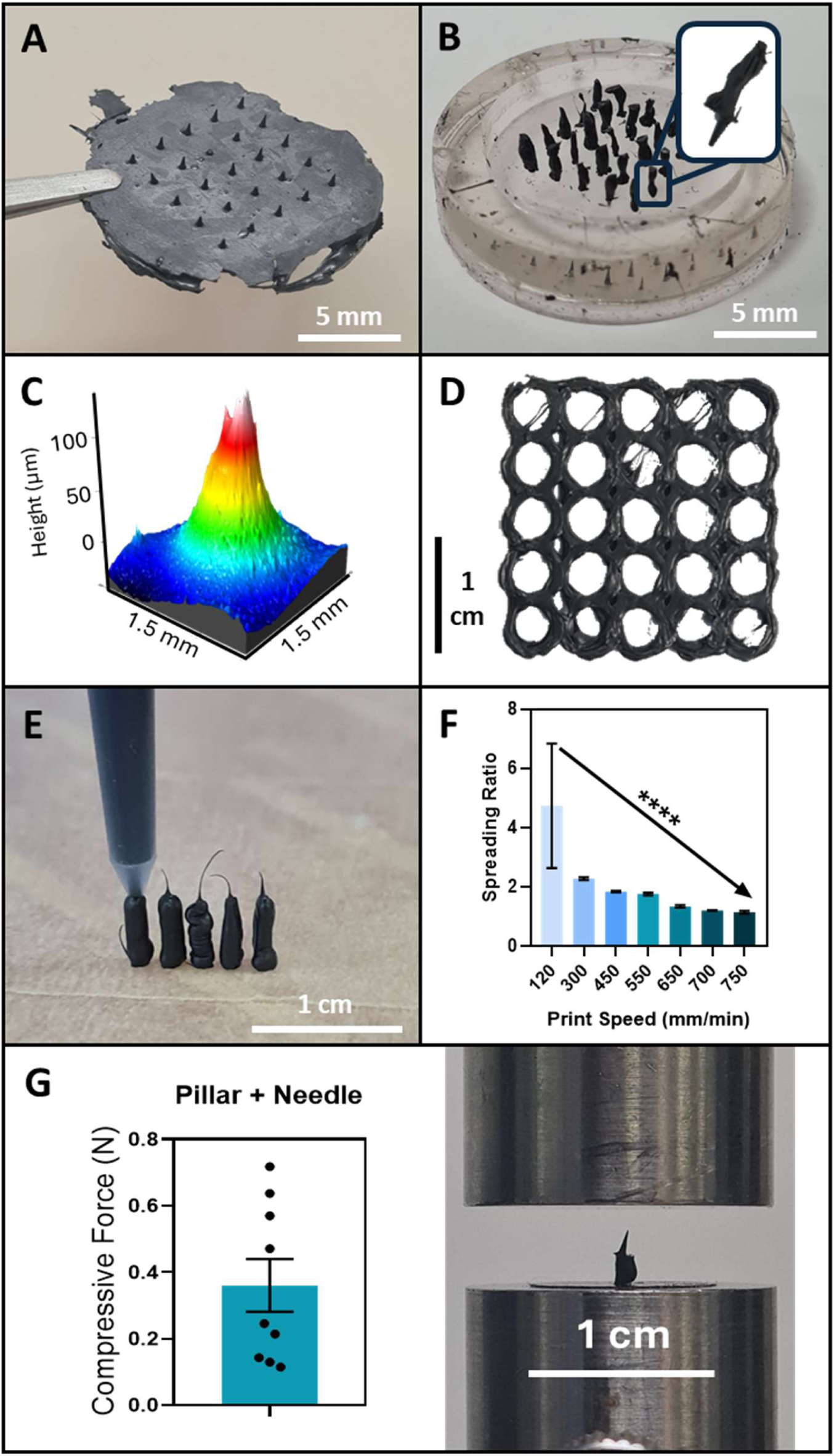
3D printing of PolyGraph: **A)** Dry-cast microneedle array. **B)** Microneedle array with pillar backing, in silicone mold. **C)** White-light interferometry profile of an individual microneedle. **D)** Cylindrical scaffold 3D printed from PolyGraph10% for tissue engineering applications. **E)** Optical images of 3D-printed pillars, enabling microneedle isolation. **F)** Determination of optimal printing speed for reduced spreading ratio in solvent 3D printing of PolyGraph. **G)** Compressive modulus testing of microneedles, assessing robustness for tissue insertion. Scale bars: A-B, 5 mm; D, E, G, 1 cm. Significances: ****p < 0.0001.

**Figure S14.**
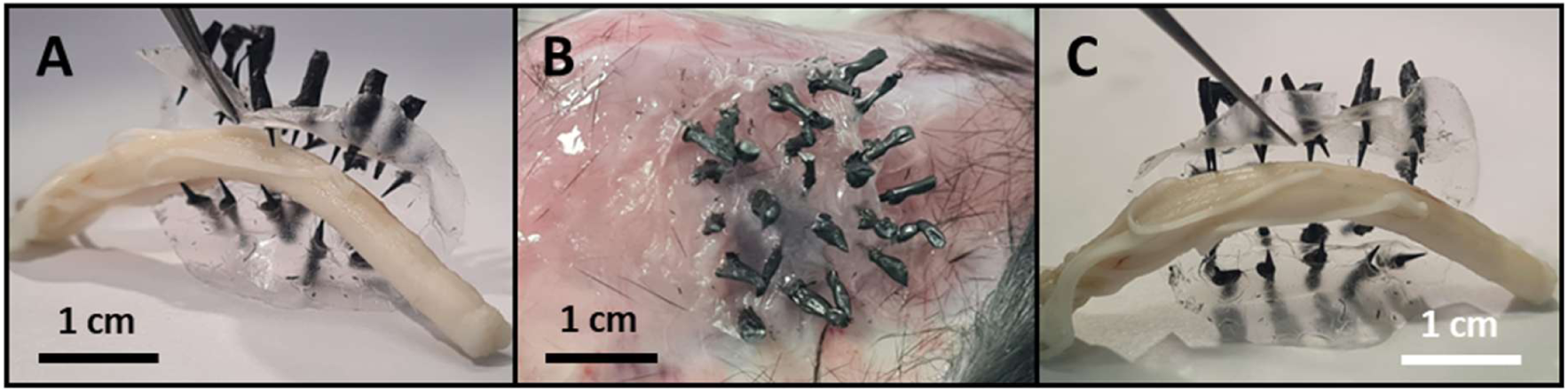
Ex-vivo application of flexible microneedle arrays: A-C) Images of flexible PolyGraph microneedle arrays on ex-vivo samples of embalmed rat spinal cord (A & C), and mouse back tissue (B), demonstrating conformability and penetration. Scale bars 1 cm.

**Figure S15.**
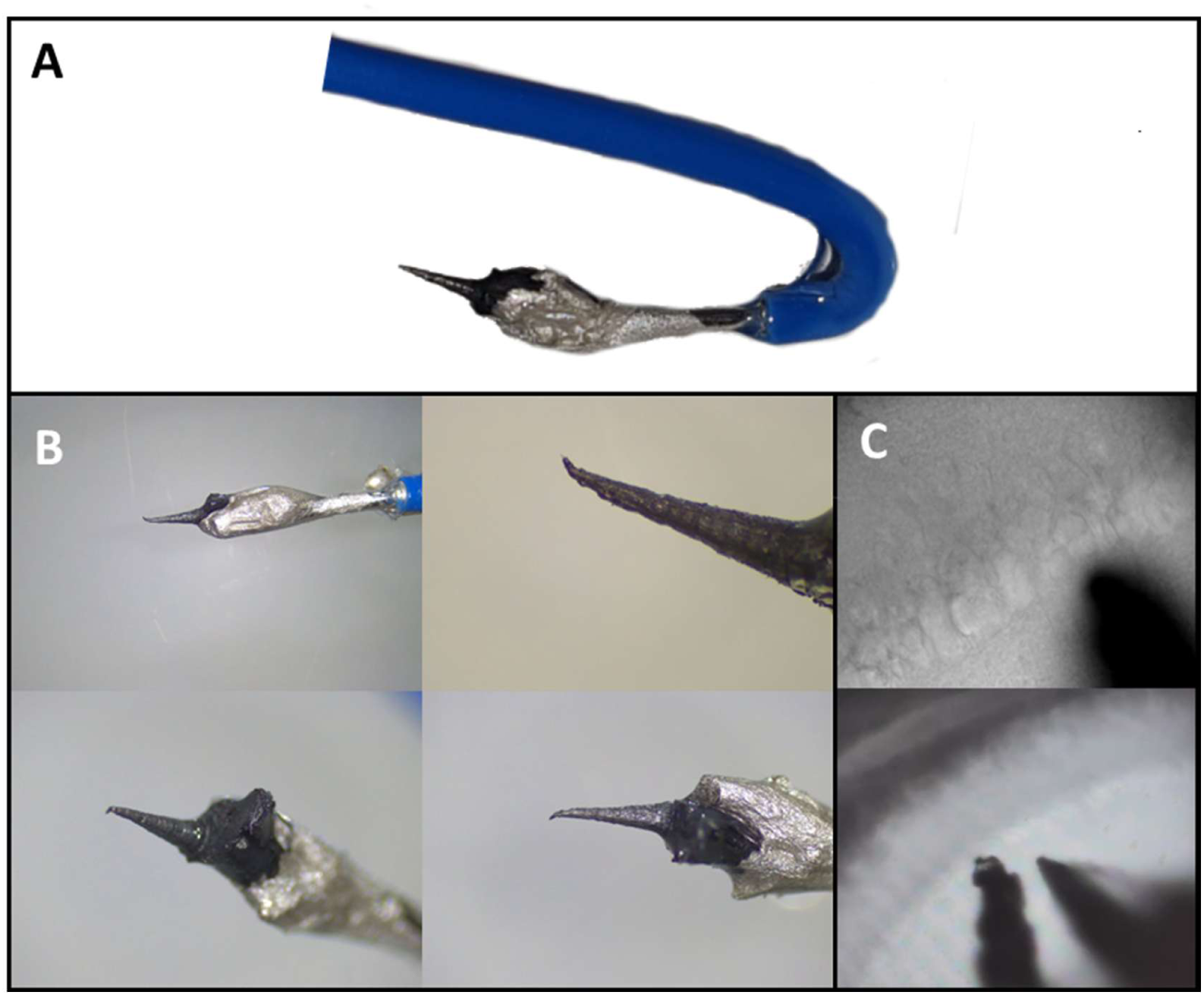
PolyGraph microneedles prepared for electrophysiological measurements: A-B) PDMS-sheathed PolyGraph microneedle electrodes following NaOH and AuPd treatments, connected to wiring using silver paint. **C)** Optical microscope image of microneedles inserted into stratum radiatum of CA1 hippocampal region, showing the sharp microneedle tip surrounded by pyramidal neurons.

**Figure S16.**
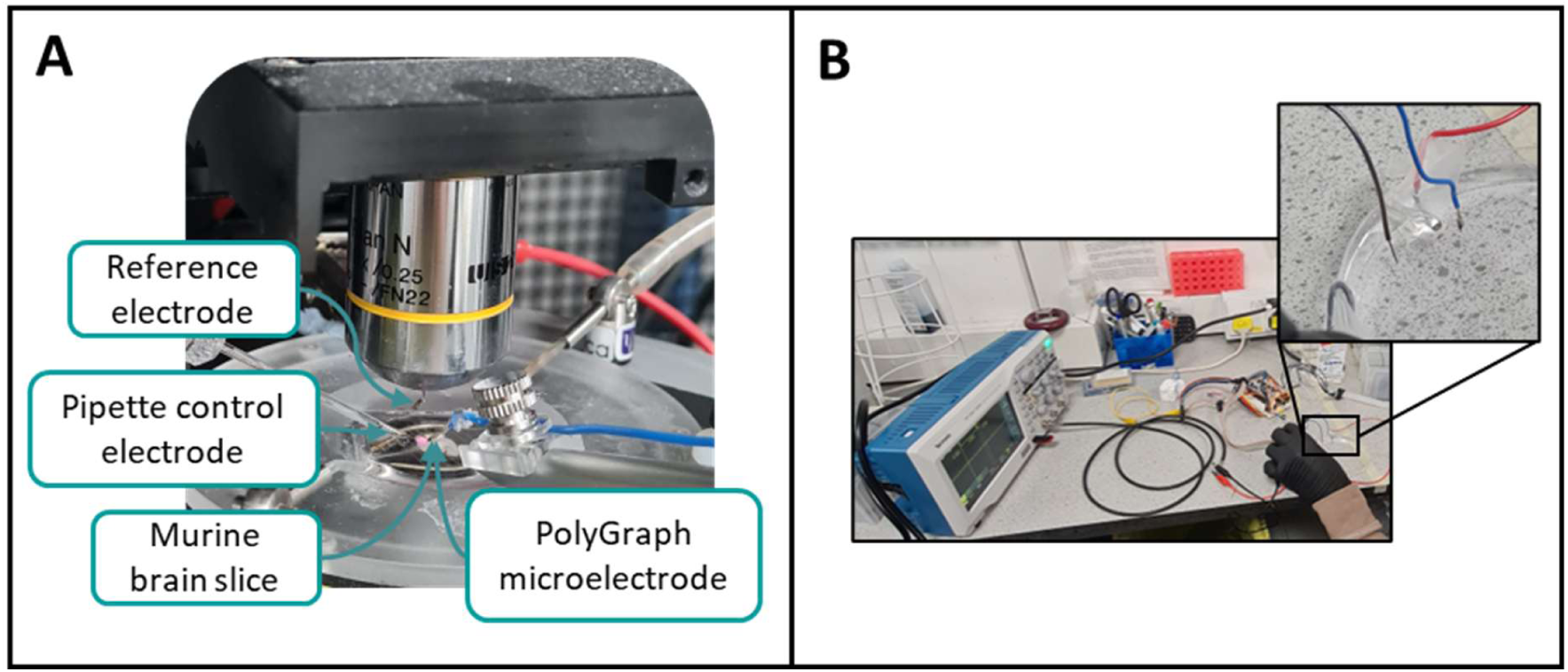
Electrophysiological recording and stimulation with PolyGraph microneedles: **A)** Experimental setup for recording local field potentials from murine brain slices, showing reference, control pipette, and PolyGraph microelectrodes. **B)** Stimulation setup with oscilloscope capturing waveform delivered through PolyGraph microneedle.

**Figure S17.**
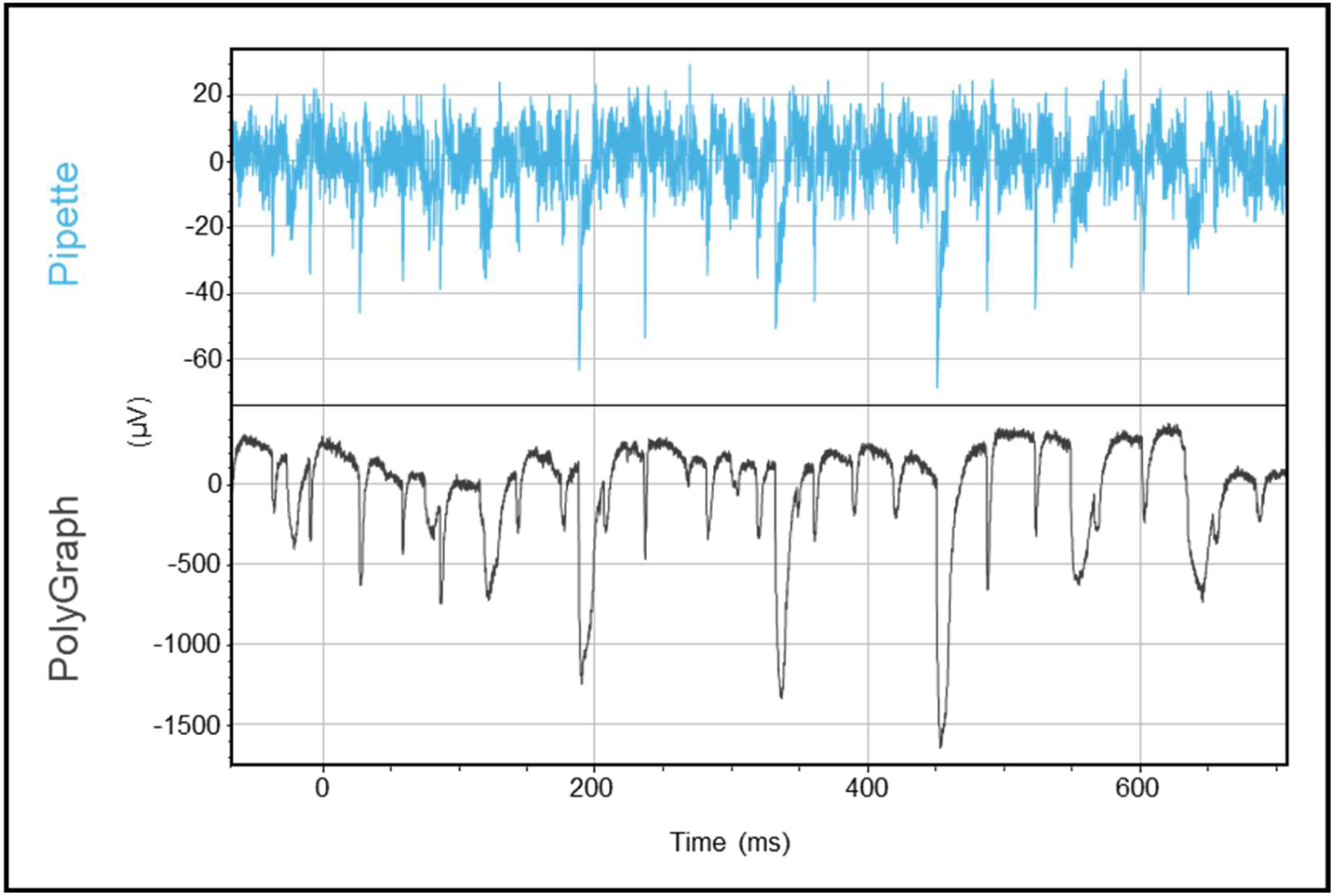
Representative raw electrophysiological trace for PolyGraph & pipette electrodes: Raw trace, showcasing increased signal-to-noise ratio (SNR) and high correlation of event waveforms recorded with PolyGraph electrodes. Y-scaling arbitrary due to differing amplifier gains.

**Figure S18.**
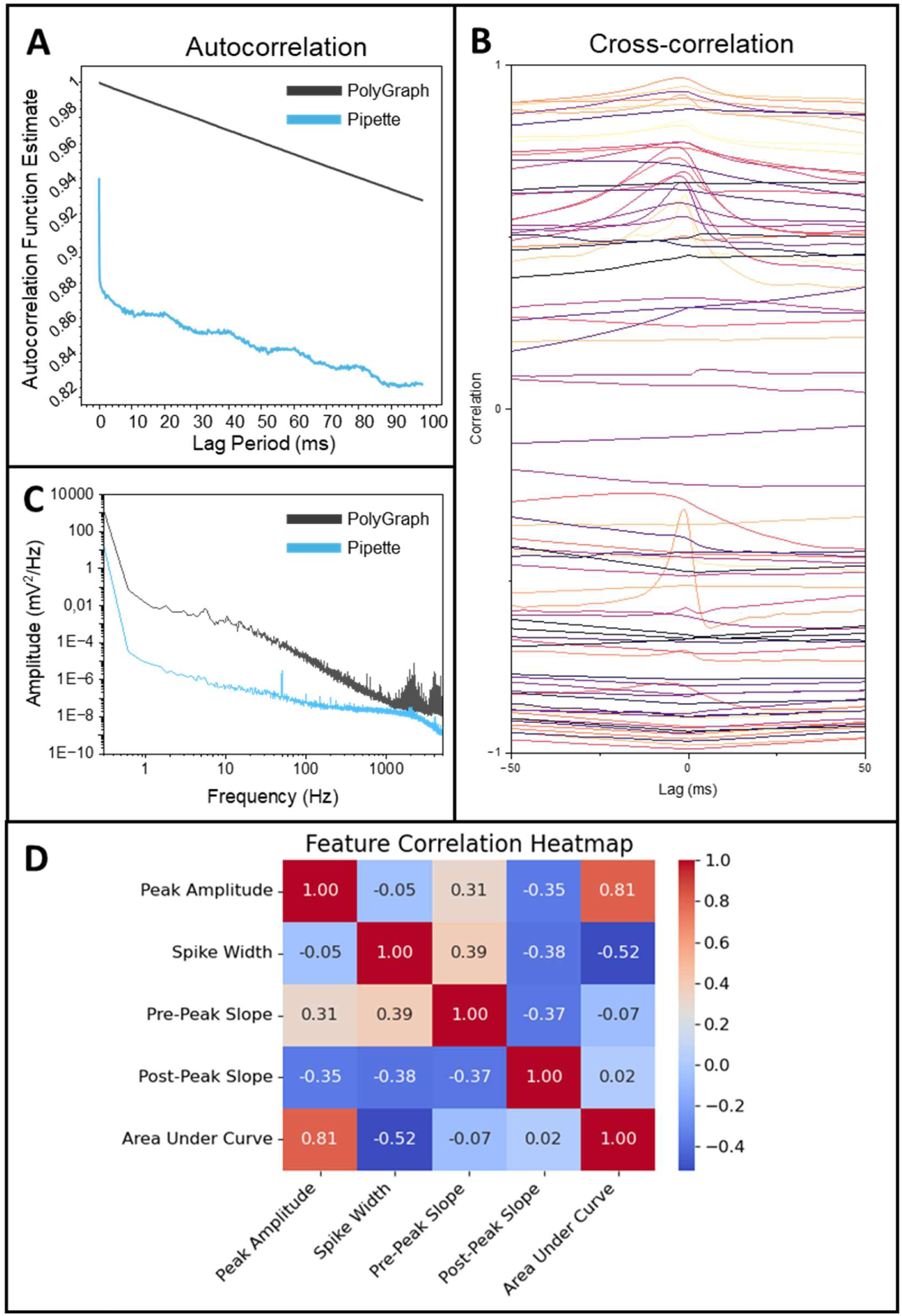
Correlation analysis of electrophysiological recordings: **A)** Autocorrelation plot, indicating the temporal coherence of PolyGraph and pipette electrodes, and their relationship with themselves in the time domain, indicating lower noise and higher temporal stability in the PolyGraph signal. **B)** Cross-correlation plots of matching sweeps for the PolyGraph and pipette electrodes, with strong correlation corroborating physiological origin of recorded events. **C)** Power spectra for each electrode, indicating higher power at all frequencies for the PolyGraph electrode, with positive implications for signal-to-noise ratio. **D)** Heatmap of correlations between extracted event features.

**Figure S19.**
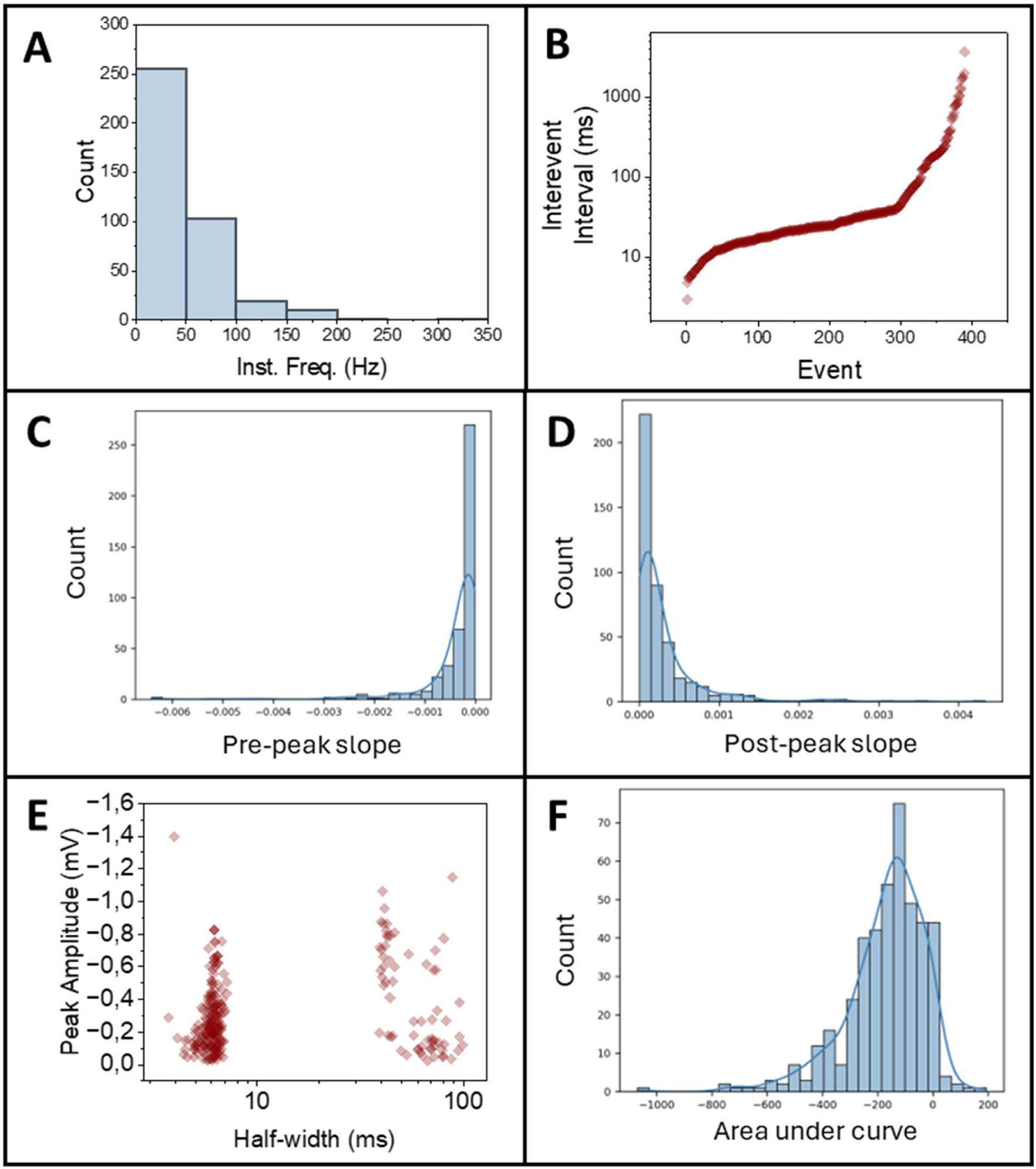
Event statistics for recordings using PolyGraph electrode: **A)** Instantaneous event frequency distribution. **B)** Interevent intervals, showing burst-like behaviour. **C-D)** Histogram of pre- (C) and post-peak (D) slopes of events. **E)** Scatter plot of peak amplitude vs half-width of events. **F)** Histogram of area under event trace, indicative of overall event magnitude.

**Figure S20.**
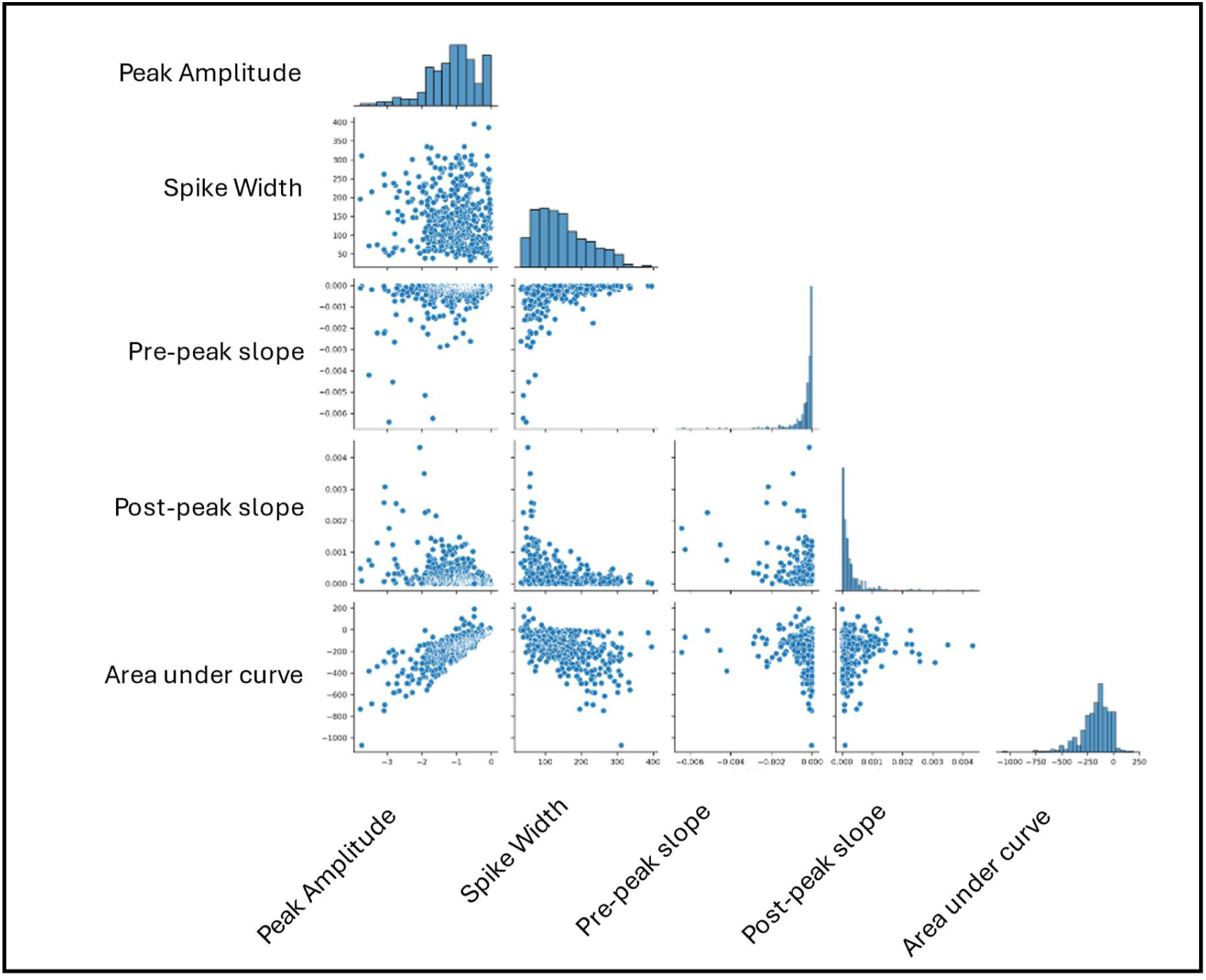
Pairwise feature analysis of events recorded by PolyGraph electrodes: Paired scatter plots and histograms of extracted event features (peak amplitude, spike width, pre- and post-peak slopes, and area under curve), showing relationships and distribution across the dataset.

**Figure S21.**
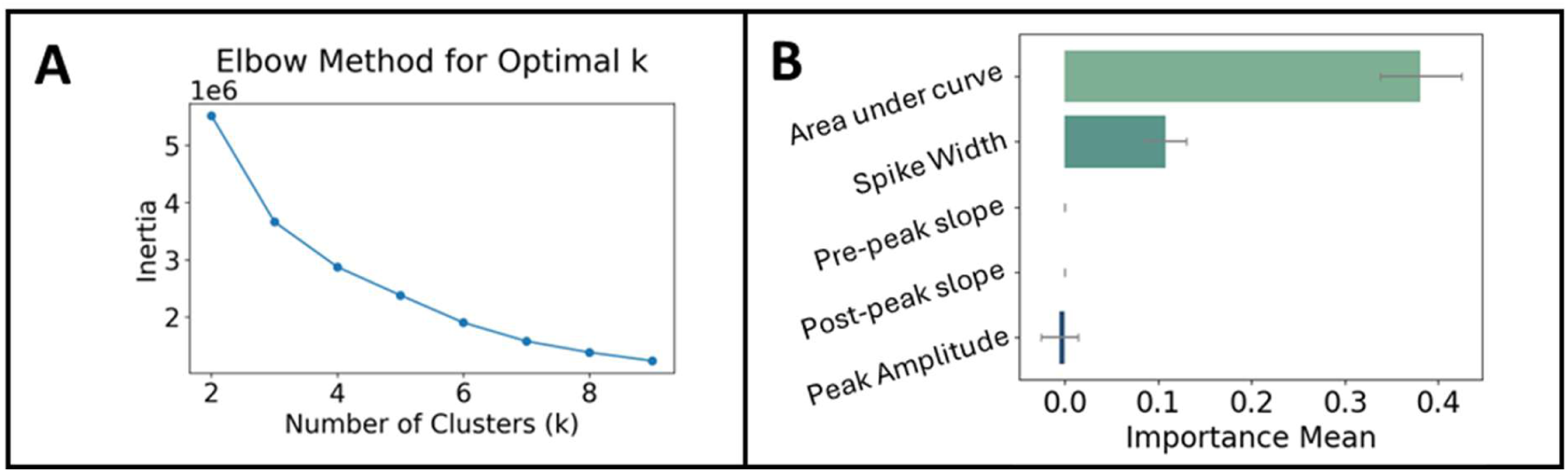
Clustering and classification of PolyGraph-recorded events: **A)** Elbow plot determining optimal number of clusters (k) for k-means clustering. **B)** Feature importance plot for MLP classifier, showing relative contribution of each feature to classification outcome.

**Figure S22.**
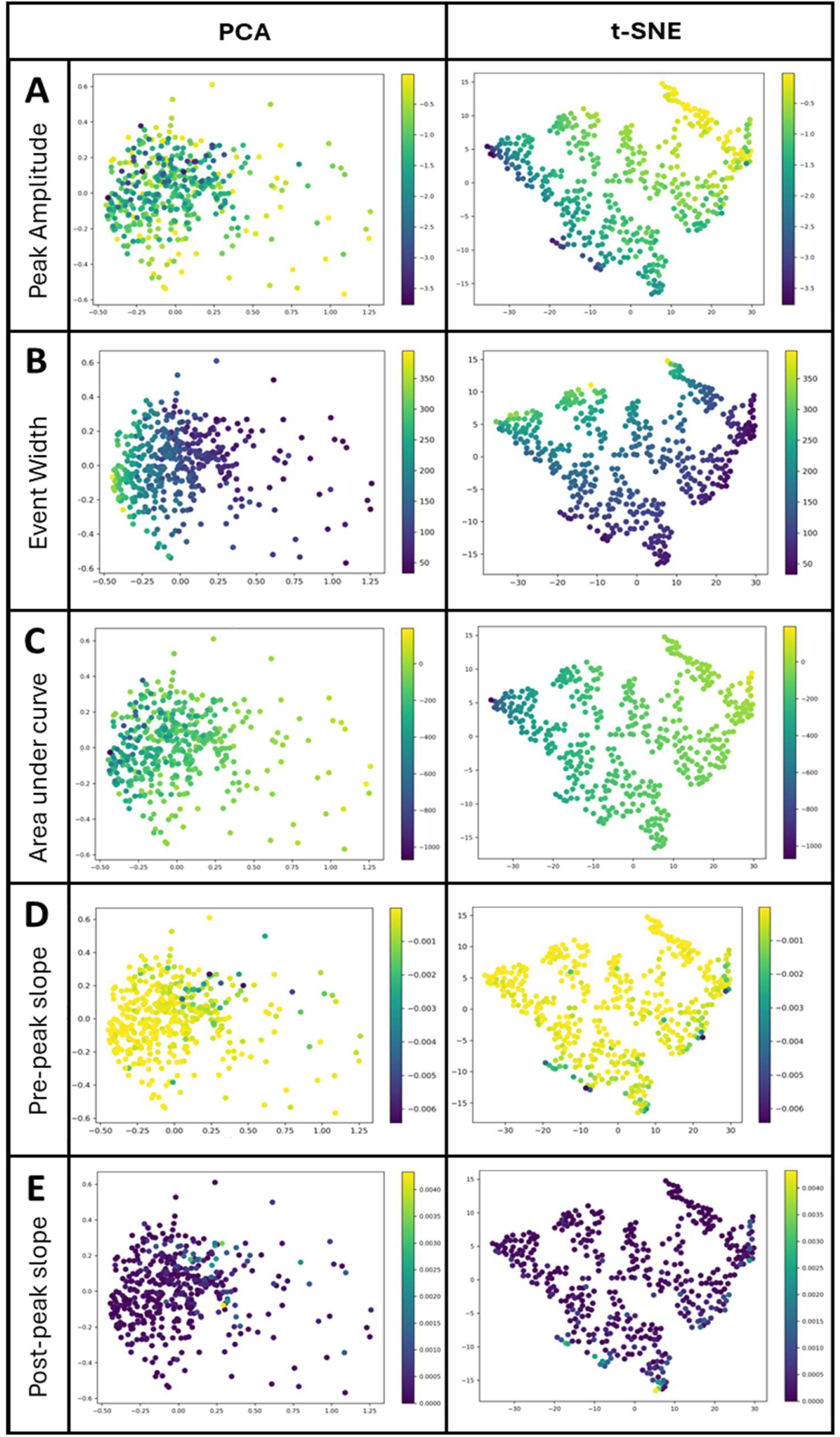
Dimensionality reduction of event features for clustering: A-E) Scatter plot results of principal component analysis (PCA, left) and t-distributed Stochastic sNeighbour Embedding (t-SNE, right) methods for feature clustering, coloured according to peak amplitude (A), event width (B), area under curve (C), pre-peak slope (D), and post-peak slope (E).

